# Olfactory receptor coexpression and co-option in the dengue mosquito

**DOI:** 10.1101/2024.08.21.608847

**Authors:** Elisha David Adavi, Vitor L. dos Anjos, Summer Kotb, Hillery C. Metz, David Tian, Zhilei Zhao, Jessica L. Zung, Noah H. Rose, Carolyn S. McBride

## Abstract

The olfactory sensory neurons of vinegar flies and mice tend to express a single ligand-specific receptor. While this ‘one neuron-one receptor’ motif has long been expected to apply broadly across insects, recent evidence suggests it may not extend to mosquitoes. We sequenced and analyzed the transcriptomes of 46,000 neurons from antennae of the dengue mosquito *Aedes aegypti* to resolve all olfactory, thermosensory, and hygrosensory neuron subtypes and identify the receptors expressed therein. We find that half of all olfactory subtypes coexpress multiple receptors. However, coexpression occurs almost exclusively among genes from the same family—among odorant receptors (ORs) or among ionotropic receptors (IRs). Coexpression of ORs with IRs is exceedingly rare. Many coexpressed receptors are recent duplicates. In other cases, the recruitment or co-option of single receptors by multiple neuron subtypes has placed these genes together in the same cells with distant paralogs. Close examination of data from *Drosophila* reveal rare cases of both phenomena, indicating that the olfactory systems of these two species are not fundamentally different, but instead fall at different locations along a continuum likely to encompass diverse insects.

## INTRODUCTION

The olfactory systems of vinegar flies and mice share a common molecular and circuit logic despite what are likely to be independent evolutionary origins^1,2^. Airborne chemicals are detected by large arrays of olfactory sensory neurons (OSNs) scattered across peripheral tissues. Each sensory neuron tends to express a single ligand-specific receptor, and all neurons that express the same receptor converge on the same spatially discrete glomerulus in the brain^2,3^. The singular expression of just one receptor per olfactory sensory neuron is often highlighted as a way to limit the tuning breadth and overlap of individual neurons/glomeruli, enabling discrimination among odorants via a combinatorial code^4,5^.

Regardless of its selective advantages, the fact that both mouse and vinegar fly OSNs canonically express just one receptor suggests that this molecular motif is functionally important and should apply broadly across vertebrates and insects alike. Yet it has been clear for over a decade that the mosquito *Aedes aegypti* does not conform. *Ae. aegypti* is a tropical mosquito that specializes in biting humans and serves as the primary vector of dengue, Zika, chikungunya and yellow fever viruses^6^. Biting females rely heavily on their sense of smell to identify humans^7^ and express about twice as many receptors in olfactory tissues (n∼130)^8,9^ as they have glomeruli in their antennal lobes (n∼60-80)^10–13^. Recent work suggests that this mismatch is at least partly explained by coexpression of multiple receptors within individual OSNs^12^. Insect olfactory receptors come from two large families, the odorant receptors (ORs) and ionotropic receptors (IRs). There is evidence that coexpression may occur among both receptors from the same family and receptors from different families (Fig. 1A).

**Figure 1.**
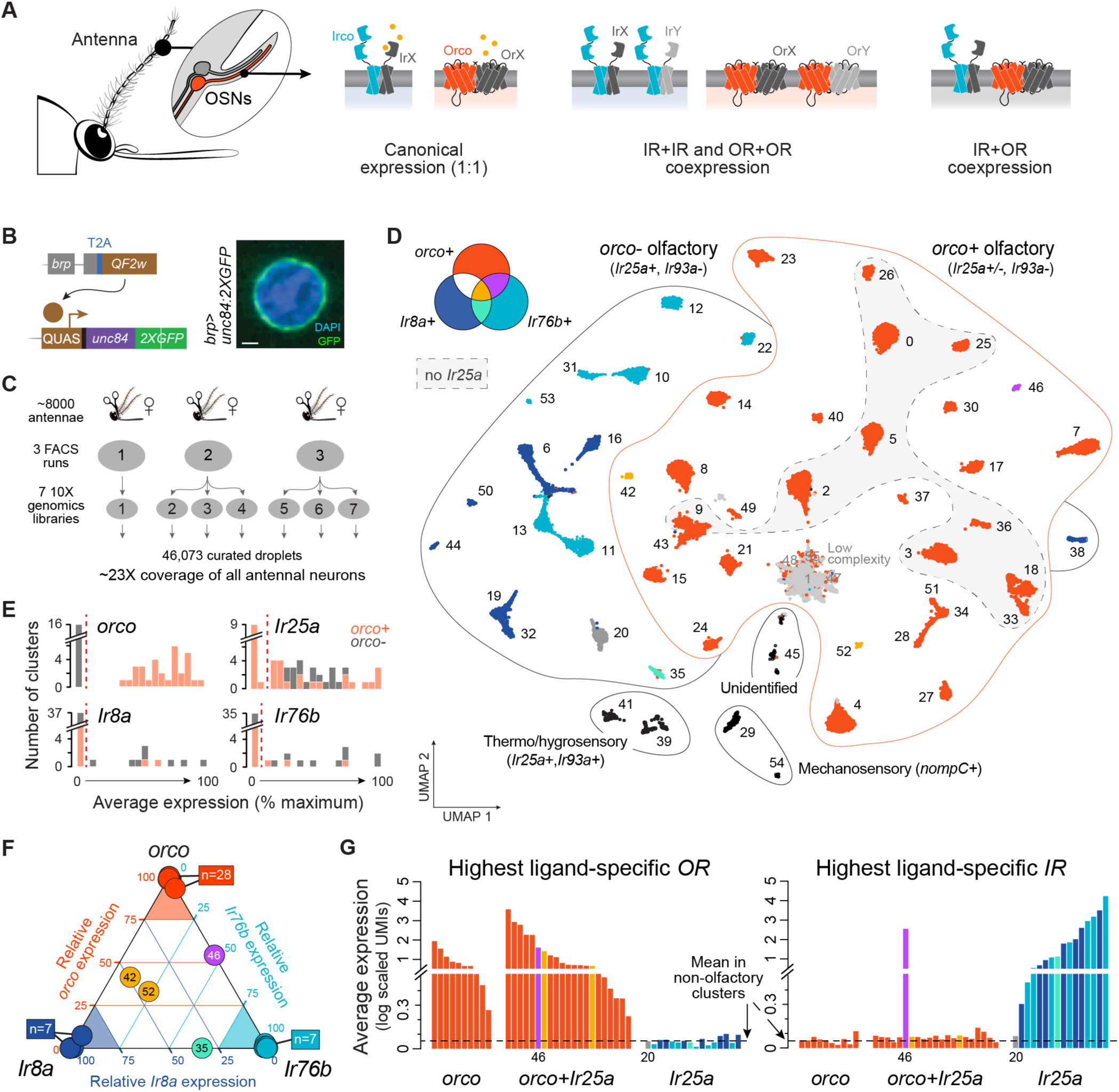
Segregation of ligand-specific *ORs* and *IRs* across female *Ae. aegypti* antennae. (**A**) Three models of receptor expression in mosquito olfactory sensory neurons (OSNs): canonical expression of one receptor per neuron (left), within-family coexpression (middle), between-family coexpression (right). Orco and ‘Irco’ stand for conserved OR and IR family coreceptors. (**B**) Transgenic strategy used to express nuclear membrane-bound GFP in neurons. Confocal image of labeled nucleus. Scale bar, 1 μm. (**C**) Single nucleus RNAseq data collection schematic. (**D**) Annotated UMAP of 46,073 nuclei from female antennal neurons (n=52 major clusters). OSN cluster colors reflect on/off expression of *orco*, *Ir8a*, and *Ir76b* as shown. (**E**) Average expression of olfactory co-receptors in OSN clusters, computed as percent gene-specific maximum. Red dashed lines show cutoffs used to call expression in (D). (**F**) Triangle plot illustrating segregated expression of *orco*, *Ir8a*, and *Ir76b*. Position of each OSN cluster (circle) reflects relative expression of the 3 genes (where expression is first computed as percent max as in (E)). (**G**) Expression level of the most highly expressed ligand-specific OR (left) or IR (right) in each OSN cluster. Cluster 46 (purple) is the only cluster that expresses substantial levels of both an OR and an IR. Cluster 20 (dark grey) expresses neither ORs nor IRs (but instead an ammonium transporter, see Fig. 3). Dashed black line shows mean for mechanosensory neurons. Colors in (F–G) as in (D).

Coexpression of ORs with IRs is particularly surprising. The neurons that express these two types of receptors were originally thought to be segregated in developmentally and anatomically distinct sensory hairs and to make up two independent olfactory subsystems^14,15^. However, a conserved coreceptor for the IR family was recently shown to be expressed in a large fraction of OR neurons in both *Drosophila*^16^ and *Ae. aegypti*^12^, generating excitement about the possibility that ligand-specific receptors from the two families also enjoy broad coexpression. Indeed, there are hints that both within- and between-family coexpression are widespread in *Ae. aegypti*, but the best data come from a small subset of OSN subtypes on the maxillary palp^12^. The full extent and nature of coexpression in this important insect remain fuzzy.

Here we conduct single-nucleus RNA sequencing of 46K neurons from female antennae to generate a comprehensive map of neuronal diversity and expression across the primary olfactory organ of *Ae. aegypti*. We find that coexpression is common among ligand-specific receptors from the same family but that coexpression between *ORs* and *IRs* is extremely rare, contrary to recent expectations. While many examples of within-family coexpression involve tandem duplicates, we also identify an unusual subset of *ORs* that have been co-opted by multiple OSN subtypes and are now expressed side-by-side with distant paralogs. Our work provides a new perspective on olfactory organization in insects and specific information about a mosquito that uses its sense of smell to transmit dangerous arboviruses to hundreds of millions of people each year.

## RESULTS

### Segregated expression of ligand-specific ORs and IRs across antennal neurons

We used single-nucleus RNA sequencing to construct a comprehensive molecular atlas of neurons from adult female antennae. Antennae harbor nearly all of the 60+ adult OSN subtypes (excluding only 3 housed on the maxillary palp) and therefore provide a broad picture of OSN diversity and organization. We first developed a transgenic strategy to label the nuclear envelope of all neurons with GFP (Fig. 1B)^17,18^ so that they could be sorted from other dissociated antennal nuclei and sequenced efficiently (Fig. 1C, Fig. S1). We then optimized *in silico* mRNA signal detection by updating the AaegL5 genome annotation in two important ways; we added a handful of missing ORs, IRs, and GRs and systematically extended 3’ UTRs where necessary to capture read pileups falling within 750bp of the annotated transcription stop site of any gene (Fig. S2, see Methods). Subsequent preprocessing, ambient RNA decontamination, and doublet removal (Fig. S3) left us with data from a total of 46,073 curated droplets, hereafter referred to as ‘nuclei’ or ‘neurons’. These data represent 23X coverage of the ∼2000 neurons present on a single female *Ae. aegypti* antenna^19^.

Initial analysis of the full dataset using the UMAP algorithm resulted in 52 well defined clusters (Fig. 1D, Fig. S4–5). We identified two clusters of putative mechanosensory neurons expressing the mechanotransduction channel *nompC*^20^, two clusters of putative heat- and humidity-sensing neurons expressing the non-olfactory IR coreceptor *Ir93a*^21^, and one cluster of unidentified neurons (Fig. 1D, Fig. S5). The remaining 47 clusters were classified as OSNs due to expression of one or more conserved olfactory coreceptors (see below). While some of these clusters are likely heterogenous, representing multiple related OSN subtypes, they allow an initial look at gross patterns of OR and IR expression.

We identified discrete expression thresholds for each olfactory coreceptor (Fig. 1E) and examined patterns of overlap across OSN clusters. Ligand-specific ORs work in complex with the *OR* coreceptor *orco*^14,22^, which showed a striking on/off pattern of expression leading to a primary classification of OSN clusters as *orco-* or *orco+* (Fig. 1D–E). Ligand-specific IRs coreceptors, *Ir25a*, *Ir8a*, and *Ir76b*^14,23,24^. We observed minimal expression of *Ir8a* and *Ir76b* in *orco+* clusters: only 3 of 31 *orco+* clusters expressed one or both of these genes (Fig. 1D,1F). In contrast, *Ir25a* was broadly coexpressed with *orco*. It was present in 21 of 31 *orco+* clusters as well as all *orco-* OSN clusters (Fig. 1D).

The presence of *Ir25a* in over half of *orco+* clusters corroborates recent findings^12,16^ and suggests that these unrelated receptor families may collaborate to define the odor tuning of many OSNs. If true, we should see coexpression of not only coreceptors, but also ligand-specific *ORs* and *IRs*. Strikingly, however, only 1 of 21 *orco+*/*Ir25a+* clusters expressed ligand-specific receptors from both families at appreciable levels (Fig. 1G, #46). The remaining 20 clusters expressed at least one *OR* (Fig. 1G, left), but contained *IR* transcripts at background levels, similar to those observed in *orco*-only or non-olfactory neurons (Fig. 1G, right). While we cannot rule out the possibility that even very low levels of *IR* expression mediate the production of functional IRco/IR complexes, we instead propose that *Ir25a* has a non-odor-tuning function in the vast majority of *orco+* OSNs in which it is found and see no reason to abandon the longstanding view of two largely distinct olfactory subsystems in insects.

### Twelve subtypes of heat- and humidity-sensing neurons

Before further exploring olfactory neurons, we sought to test the ability of our data to resolve rare cell types by focusing on thermo- and hygrosensory neurons (THSNs). *Ae. aegypti* antennae harbor at least 4 types of THSNs, termed heating, cooling, dry, and moist cells^19,21,25^. Most of these cells express the non-olfactory IR coreceptor *Ir93a*^21^, and research in both *Drosophila* and mosquitoes suggests that *Ir93a* works in concert with modality-specific IR partners, including *Ir21a* (cooling), *Ir40a* (dry), and *Ir68a* (moist)^21,26–28^. Importantly, THSNs are sparsely distributed across the most basal and distal antennal segments (Fig. 2A), with some subtypes likely comprising just 1-3 neurons per antenna^19,21,25^.

**Figure 2.**
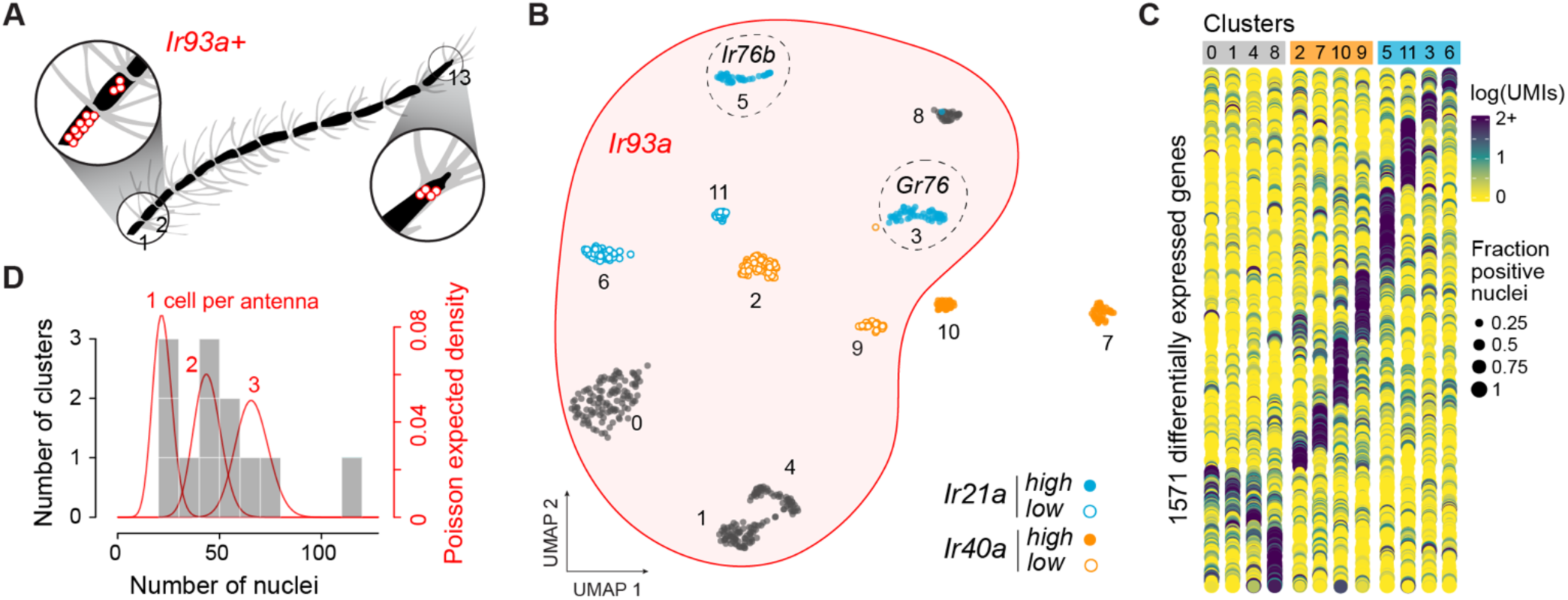
Deep snRNA-sequencing resolves twelve small subpopulations of candidate thermo- and hygrosensors. (**A**) Location of *Ir93a+* thermo- and hygrosensory neurons on segments 1, 2, and 13 of female *Ae. aegypti* antennae^19,21^. (**B**) Annotated UMAP showing reclustering of 589 nuclei from two *Ir93a+* clusters in the original analysis (Fig. 1D). Note that 2 of the 12 new clusters do not expres *Ir93a*. (**C**) Dotplot showing expression of 1571 genes differentially expressed across clusters (log2FC>0.3). (**D**) Histogram showing number of nuclei in each cluster (grey bars) overlaid by the expected densities for cell types comprising 1, 2, or 3 neurons per antenna given 23X coverage (red lines).

*In silico* subsetting and reclustering of nuclei from the two *Ir93a+* clusters in the all-neuron analysis revealed 12 putative subtypes of THSNs (Fig. 2B, Fig. S6–7), each with a strikingly unique transcriptional profile (Fig. 2C).

The cooling receptor *Ir21a* was expressed in four clusters, one of which was notable for coexpression of the IR coreceptor *Ir76b*, and another for coexpression of a gustatory receptor (Fig. 2B, Fig. S7), patterns that have not been documented in *Drosophila*. The dry receptor *Ir40a* was also expressed in four clusters, two of which did not express *Ir93a* despite having been lumped with other *Ir93a+* cell types in the all-neuron analysis (Fig. 2B, Fig. S7). These *Ir40+*/*Ir93a-* nuclei may correspond to elusive heating cells^21^. As predicted, the number of nuclei in all but one of the 12 THSN clusters ranged from 25-75, consistent with cell types that comprise just 1-3 cells per antenna (Fig. 2D, Fig. S8). Taken together, this analysis reveals exciting cellular diversity among THSNs, along with candidate receptors and a list of marker genes that may be used to gain genetic access for functional studies. It also confirms the ability of our data to resolve rare cell types.

### *IR* coexpression across *orco-* OSNs

Coexpression of ligand-specific *ORs* and *IRs* is rare (Fig. 1G), but coexpression among receptors from the same family may still be widespread. We first set out to explore this possibility among *IRs* in *orco-* OSNs. To resolve as many neuron subtypes as possible, we subsetted and iteratively cleaned and reclustered nuclei from the 16 original *orco-* clusters (Fig. 1D, see Methods). This process identified 4 additional clusters or subclusters (hereafter simply ‘clusters’) for a total of 20 (Fig. 3A, Fig. S6,S9–11). As seen in the original analysis, all clusters expressed *Ir25a*, and almost all additionally expressed either *Ir8a* or *Ir76b*, but rarely both (Fig. 3A–B). The total number of *orco-*, *Ir8a+*, and Ir76b+ clusters in our analyses agreed remarkably well with the number of antennal glomeruli identified in each category in recent work (Fig. S9B)^12^. We therefore conclude that most, if not all, clusters represent single, homogenous OSN subtypes with unique glomerular targets.

**Figure 3.**
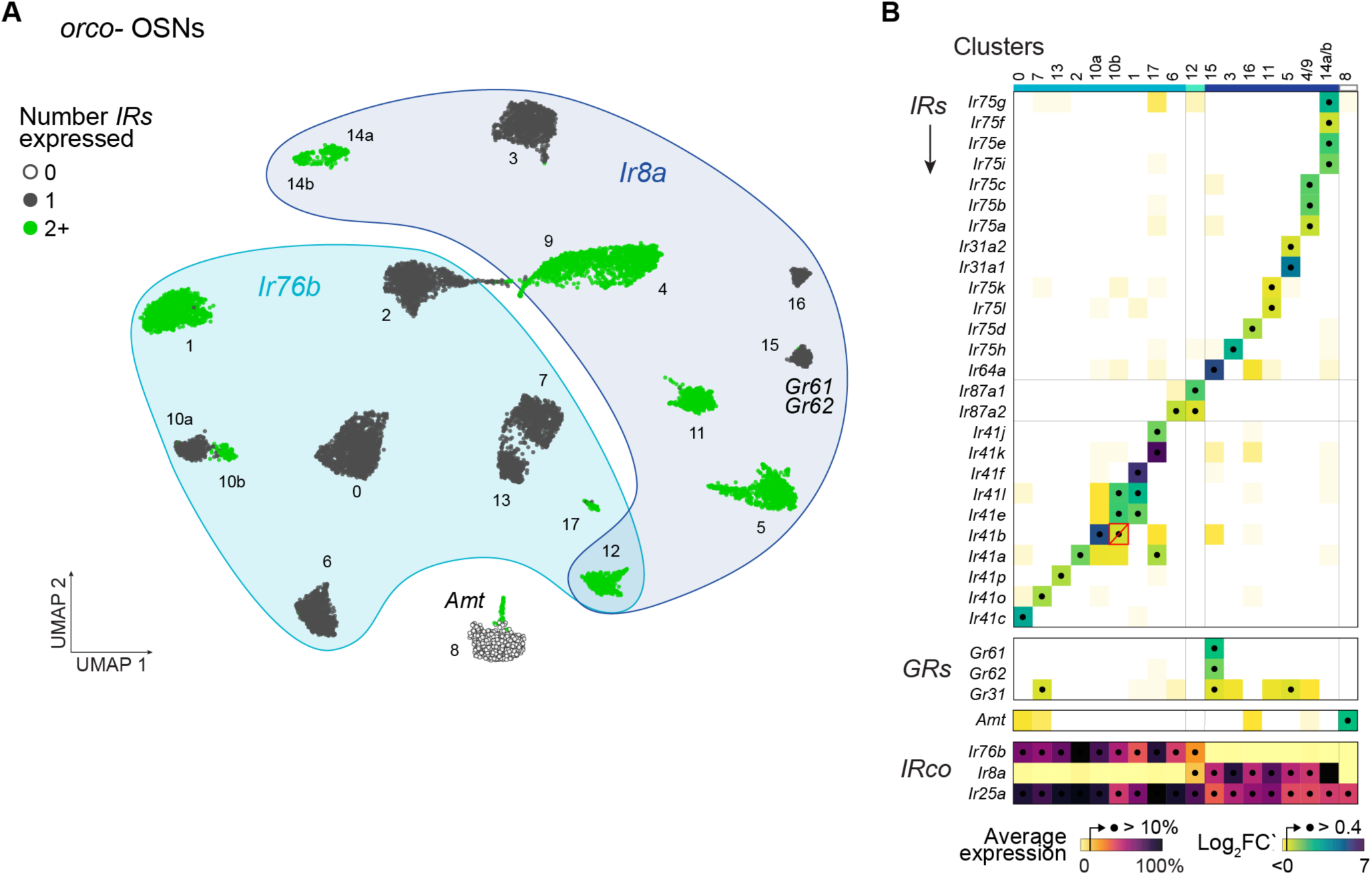
IR coexpression across *orco-* olfactory neurons. (**A**) Annotated UMAP showing reclustering of 12,243 nuclei assigned to *orco-* OSN clusters in the original analysis (Fig. 1D). (**B**) Differential expression of ligand-specific receptors across clusters. All *IRs, ORs*, and *GRs*, exceeding a log2FC’ cutoff of 0.4 in any cluster are shown, but few *GRs* and no *ORs* met this criterion. The red box/slash marks an expression call (*Ir41b* in cluster 10b) that is likely spurious. Clusters expressing the same receptors were merged (#4/9, 14a/b). See Fig. S12 for log2FC’ and log average expression in all clusters examined separately. *IR* co-receptor expression shown at bottom as percent gene-specific maximum.

To identify ligand-specific receptors expressed in each *orco-* subtype, we looked for *IRs* that were differentially expressed across clusters (Fig. 3B). More specifically, we computed log2FC’ for all *IRs* in all clusters and chose a cutoff (0.4) that separated a large number of near-zero values from a much smaller number of clearly elevated values (Fig. S12A). Any *IR* exceeding this threshold was considered expressed in the given cluster. Use of an alternative absolute expression cutoff produced nearly identical expression calls (Fig. S12B–C). We also tested *ORs* and *GRs*, but observed no expression of the former and only sporadic expression of the latter (Fig. 3B).

Nine of the 20 putative *orco-* OSN subtypes expressed a single ligand-specific *IR*, while 10 expressed two or more *IRs* (Fig. 3A–B). Importantly, when two *IRs* were called as expressed in the same cluster, they also showed correlated expression at the neuron level (Fig. S12D–E), corroborating that these represent true cases of coexpression. The only exception involved *Ir41b*, whose elevated expression in cluster 10b likely results from a clustering artifact (Fig. 3B, red outline/slash; see also Fig. S12E). In two cases, transcriptionally similar clusters appeared to express the same or very similar subsets of *IRs* (Fig. S10, S12B–C, #4/9 and 14a/b). We conservatively assume these correspond to single OSN subtypes and show merged data in Fig. 3B.

One putative *orco-* OSN subtype expressed neither *Ir8a*, *Ir76b*, nor any ligand-specific *IR* (Fig. 3A–B, #8). Instead, this subtype expressed the ammonium transporter *Amt* (Fig. 3A–B; Fig. S11D), which is required for ammonia detection in *Drosophila*^29^, though perhaps not in malaria mosquitoes^30,31^. These neurons should be of great interest in *Ae. aegypti* as ammonia is a key host odorant, synergizing with lactic acid and carbon dioxide to attract biting females^32^.

### *OR* coexpression and multi-expression across *orco+* OSNs

We next sought to characterize patterns of *OR* expression among *orco+* sensory neurons using a similar approach—by iteratively cleaning and reclustering nuclei from the original *orco+* clusters (Fig. 1D) and using differential and absolute expression to identify ligand-specific receptors expressed therein. We identified 40 *orco+* clusters in total (Fig. 4A, Fig. S9,S13-14), which conservatively correspond to 37 putative OSN subtypes after merging pairs of clusters that expressed the same or similar receptors (Fig. S15). This number is again consistent with a 1:1 match between clusters and neural subtypes based on recent counts of *orco+* antennal lobe glomeruli (Fig. S9B)^12^.

**Figure 4.**
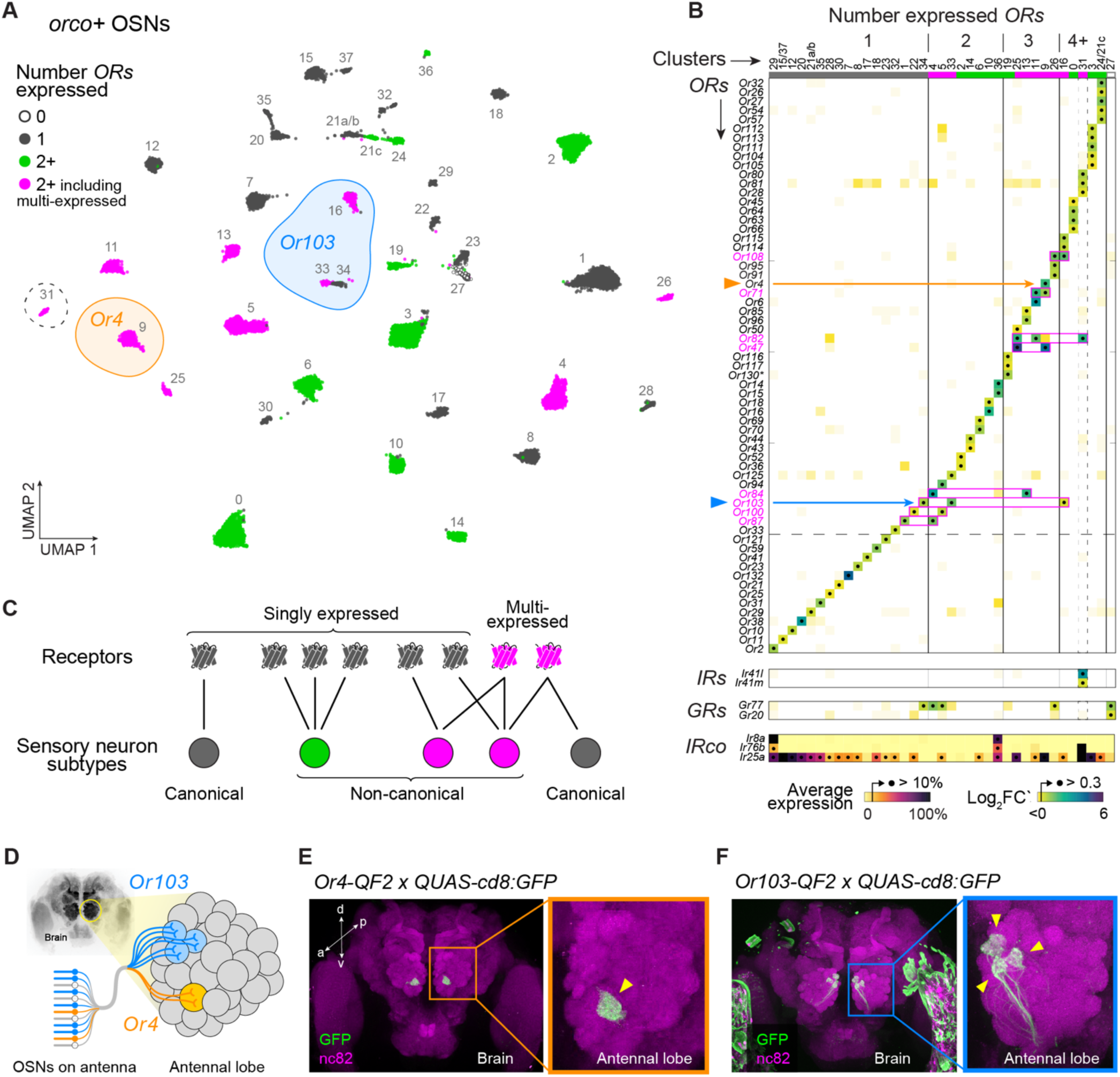
OR coexpression and multi-expression across *orco+* olfactory neurons. (**A**) Annotated UMAP showing reclustering of 28,807 nuclei assigned to *orco+* OSN clusters in the original analysis (Fig. 1D). Dashed circle marks the only cluster (#31) that expressed ligand-specific *IRs*. (**B**) Differential expression of ligand-specific receptors across 37 major clusters. All *ORs*, *IRs*, and *GRs* with log2FC’ > 0.3 in any cluster are shown (see Fig. S15 for lower cutoff). Clusters expressing the same receptors were merged (#15/37, 21a/b, 24/21c). See Fig. S15 for log2FC’ and log average expression in all clusters examined separately. *IR* co-receptor expression shown at bottom as percent gene-specific maximum. Pink boxes highlight expression calls for multi-expressed *ORs*. (**C**) Schematic illustrating classification of receptors and OSN subtypes based on patterns of expression. (**D**) OSNs (bottom left) project axons to antennal lobe glomeruli (right) according to subtype. Data in (B) suggest that *Or4* and *Or103* are expressed in 1 and 3 OSN subtypes, respectively. (**E**–**F**) Antibody stainings show the GFP-expressing axons of *Or4* (E) and *Or103* (F) neurons in the adult female antennal lobe. Yellow arrowheads mark target glomeruli. See Methods for details.

As seen in the original analysis, approximately two thirds of *orco+* subtypes expressed the IR coreceptor *Ir25a*, but only 1 expressed ligand-specific *IRs* (Fig. 4B, #31). In contrast, almost all *orco+* subtypes expressed at least one *OR*, and approximately half coexpressed multiple *ORs* (Fig. 4B, Fig. S15A–B). Without exception, coexpression calls were again supported by elevated correlations at the neuron level (Fig. S15D–E). *OR*+*OR* coexpression in *orco+* OSNs thus appears to be just as common as *IR*+*IR* coexpression in *orco-* OSNs.

We were intrigued to find not only coexpression, but also a phenomenon we call ‘multi-expression’. This refers to cases where the same receptor is present in multiple neuron subtypes (Fig. 4C). For example, *Or103* was expressed in three putative subtypes (Fig. 4A). It was alone in cluster 34, coexpressed with *Or125* in cluster 33, and coexpressed with *Or108*, *Or114*, and *Or115* in cluster 16 (Fig. 4B). Likewise, while *Or4* was only expressed in a single cluster (Fig. 4A, cluster 9), it was coexpressed in that cluster with two receptors (*Or47* and *Or71*) that were each present alongside different partners in second clusters (Fig. 4B). In total, our data indicate that 8 of 63 antennal *ORs* are multi-expressed (Fig. 4B) and that 13 of 37 orco+ subtypes express at least one of these genes.

To provide independent support for multi-expression, we used CRISPR/Cas9 to generate driver lines for both *Or4* and *Or103*, crossed these drivers to an existing GFP effector^33^, and counted the number of labeled glomeruli.

Importantly, our driver lines leverage the *cis*-regulatory elements of the target receptors in situ (Fig. S16), which is expected to more faithfully replicate endogenous patterns of expression than alternative approaches. If, as our snRNAseq data suggest, *Or4* is expressed in one OSN subtype and *Or103* is expressed in three OSN subtypes, then the *Or4* and *Or103* lines should drive expression of a GFP effector in neurons that target 1 and 3 glomeruli, respectively (Fig. 4D). This is exactly what we observed (Fig. 4E-F).

### Two drivers of olfactory receptor coexpression

Receptor coexpression has multiple potential sources. The most likely is recent tandem gene duplication, wherein errors in DNA replication or recombination generate two adjacent copies of an ancestral gene. New duplicates are typically coexpressed because they share or inherit the same regulatory elements^34^. Importantly, this type of gene duplication is expected to drive coexpression among genes that are both physically and phylogenetically close. A second potential source of receptor coexpression is the recruitment of preexisting receptors to novel OSN subtypes. For example, the multi-expression observed for several *ORs* may arise when a singly expressed gene is co-opted by a second neural subtype following *cis*-regulatory evolution, *trans*-regulatory evolution, or cell type birth. This process would lead to coexpression between the newly recruited receptor and other receptors in the second cell type, which we would not necessarily expect to be closely related or physically adjacent. We can think of these two phenomena as coexpression ‘by descent’ and coexpression ‘by co-option’.

To identify the source of coexpression among *Ae. aegypti IRs* and *ORs*, we examined pairwise phylogenetic and chromosomal distances for receptors that were or were not coexpressed. Strikingly, all 20 pairs of coexpressed *IRs* were closely related and located in the same genomic cluster, consistent with a model of coexpression by descent (Fig. 5A). Many coexpressed *ORs* were also closely related and physically near—especially those that were singly expressed (green dots/density in Fig. 5B). However, multi-expressed *ORs* were rarely located in the same genomic cluster as their coexpressed partners and could be either closely or distantly related, consistent with a model of coexpression by co-option (Fig. 5B). Taken together, these patterns strongly suggest that *IR*+*IR* coexpression is driven predominantly by the coinheritance of ancestral regulatory elements during gene duplication, while *OR*+*OR* coexpression is driven not only by gene duplication, but also but the co-option of a subset of receptors to new neural subtypes.

**Figure 5.**
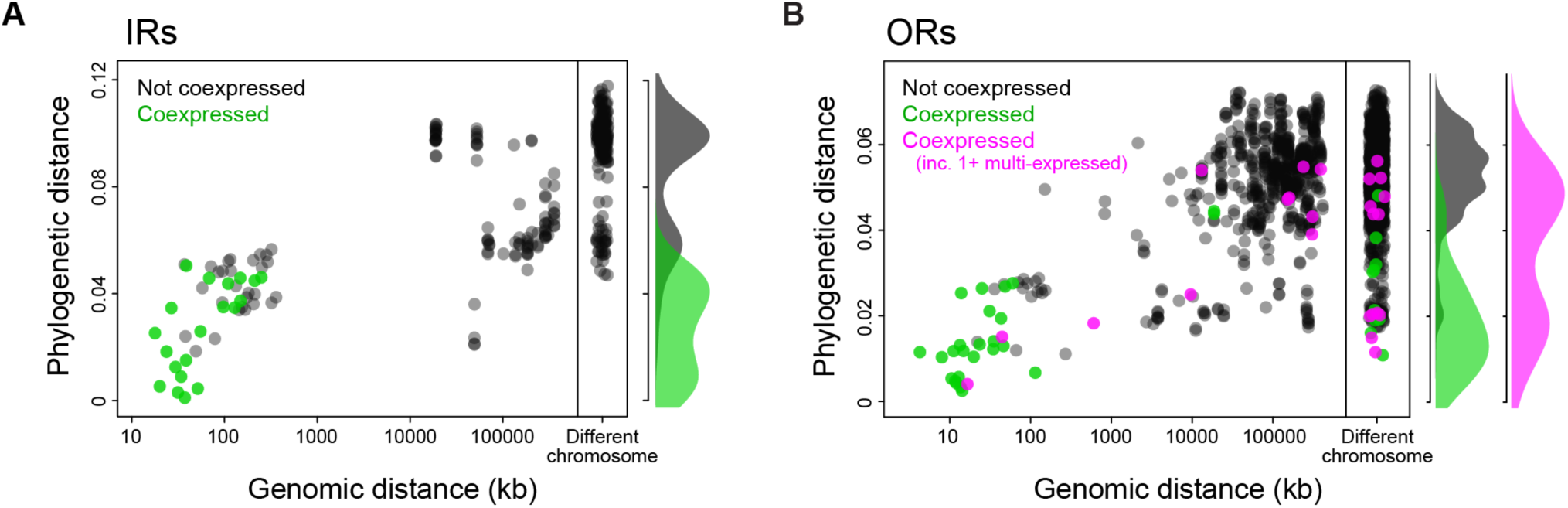
OR coexpression is driven by both receptor duplication and receptor co-option. Plots show the genomic and phylogenetic distance for pairs of ligand-specific *IRs* (A) or *ORs* (B). Color indicates whether the receptors are coexpressed in the same OSN subtype. Coexpressed *ORs* are further divided into cases where both receptors are found only in the same, single subtype (green) or at least one receptor is also found in additional subtypes (pink). Marginal densities for phylogenetic distance shown at right of each scatterplot. Phylogenetic distance in units of amino acid changes/site.

## DISCUSSION

Recent work in *Ae. aegypti* mosquitoes challenges the longstanding view that the olfactory systems of most insects will resemble those of vinegar flies and mice, with each sensory cell type expressing a single ligand-specific receptor^12^. Yet exactly how and to what extent this ‘rule’ is broken has remained uncertain. Here we use deep single-nucleus RNA sequencing data to generate a surprisingly clear picture of neuronal diversity and receptor expression across female *Ae. aegypti* antenna. We show that half of all olfactory neurons coexpress multiple receptors, but that coexpression occurs almost exclusively among genes from the same receptor family—among *IRs* or among *ORs*. Coexpression of genes from different families is rare. We also identify an unexpected evolutionary driver of coexpression—the co-option of single receptors by multiple neural subtypes.

The segregation of *ORs* and *IRs* in all but one of ∼60 antennal olfactory cell types adds nuance to the recent discovery of coexpression among coreceptors^12,16^. *Ir25a* is broadly coexpressed with *orco*, but it is unlikely to support odorant detection without a ligand-specific partner (let alone the help of a second IR coreceptor^23,24^). We instead hypothesize that *Ir25a* has an alternative, non-odor tuning function in most *orco+* cells, perhaps acting during development or regulating other aspects of sensory transduction^16^. At the same time, the rarity of OR+IR coexpression makes the few instances where it *does* occur interesting from a behavioral perspective. For example, OR+IR neurons are uniquely poised to drive innate behavioral responses to structurally unrelated compounds with the same ecological meaning^12^. It remains to be seen whether this is true in *Ae. aegypti*, but it appears likely in a recent example from female hawkmoths^35^. In addition to the one OR+IR cell we identify on antennae, ligand-specific *IRs* have been documented in two *orco*+ cells on the maxillary palp, albeit at relatively low levels^12^.

While OR+IR coexpression is rare, we confirm widespread coexpression of receptors from the same family. Inferences of coexpression based on snRNA sequencing have some inherent limitations. Of greatest concern is the possibility that some of the transcripts we detect in nuclei are not exported and/or translated. For example, recent work in the clonal raider ant revealed cotranscription of dozens of tandem ORs within single nuclei, but only transcripts from the most upstream locus made their way into the cytoplasm^36^. This interesting phenomenon is likely specific to ants, which have the largest *OR* repertoires among insects. However, other, more limited forms of post-transcriptional repression have been detected among *Drosophila* IRs^37^ and could easily be present in *Ae. aegypti*. Conversely, the three-prime sequencing used in our study and most other snRNAseq studies will miss cases of coexpression among tandem genes found on a single polycistronic transcript, as has been observed in *Anopheles* mosquitoes^38^. We do not expect these phenomena to change our overall conclusions, but further work will be needed to confirm the precise rate at which *Ae. aegypti* OSNs employ multiple ligand-specific receptors for odor detection. Moving forward it will also be critical to understand how multiple ORs or multiple IRs interact, either directly or indirectly, to define the odor tuning of an OSN.

One of our most surprising findings is that OR+OR coexpression stems not only from gene duplication, but also from the co-option of a subset of receptors by multiple olfactory cell types. We do not know exactly how such co-option occurs, but one intriguing possibility is that it sometimes coincides with the birth of new cell types. Work in *Drosophila* suggests that new olfactory neurons can evolve via the repression of programmed cell death in precursor cells that would otherwise be eliminated during development, and that these ‘undead’ neurons initially express one or more receptors already present in other olfactory cells^39^. Multi-expressed genes in *Ae. aegypti* could represent the vestiges of such a process. Regardless of exactly how, when, or why co-option occurs, multi-expressed *ORs* are clearly unique in that they are frequently coexpressed with distantly related receptors (Fig. 5B).

Our work and that of others raises the question of why and when we might expect *any* animal to express only one *vs*. multiple receptors per OSN. Singular expression is often highlighted as a way to limit the tuning breadth/overlap of individual neurons in support of combinatorial coding^4,5^. This may explain why OSNs should express *few* receptors, but it is not a satisfying explanation for why OSNs must express just *one* receptor. Strict singular expression in vertebrates likely reflects developmental constraints more than any particular coding strategy. Vertebrate olfactory systems have hundreds to thousands of OSN subtypes— perhaps too many to be efficiently specified by conventional transcription factor codes. Instead, each mouse OSN stochastically expresses a single receptor, which then defines the tuning of the neuron *and* helps direct axon development to ensure proper targeting in the brain^40^. Interestingly, there are hints that ants, which also have complex olfactory systems, have independently converged on a receptor-dependent developmental strategy that enforces singular expression^36,41^. It is harder to explain why insects with deterministic olfactory development, like *D. melanogaster*^42,43^, would so closely follow the one neuron-one receptor rule. It is possible that singular expression helps to optimize combinatorial coding, even if not strictly required, and is therefore advantageous to species that rely heavily on learning. Coexpression of small numbers of receptors, in contrast, may allow evolution to more precisely tune neurons to important resources or threats in animals for which learning is less important. Or perhaps the number of receptors expressed per neuron reflects non-ecological factors like genome size and dynamics, with selection for compact genomes favoring smaller receptor repertoires and the proliferation of repetitive elements in larger genomes facilitating receptor family expansion.

Regardless of their differences, we would argue that *Ae. aegypti* and *D. melanogaster* are fundamentally similar. *Drosophila* OSNs are more likely to express just one receptor, but exceptions exist^3,44,45^, and close examination of those exceptions reveals the same signatures of coexpression by descent and coexpression by co-option that we see in mosquitoes (Fig. 6). Patterns of receptor expression are thus qualitatively (if not quantitatively) similar. We propose that the dengue fever mosquito and vinegar fly simply lie at different positions along a continuum that is likely to encompass most insects.

**Figure 6.**
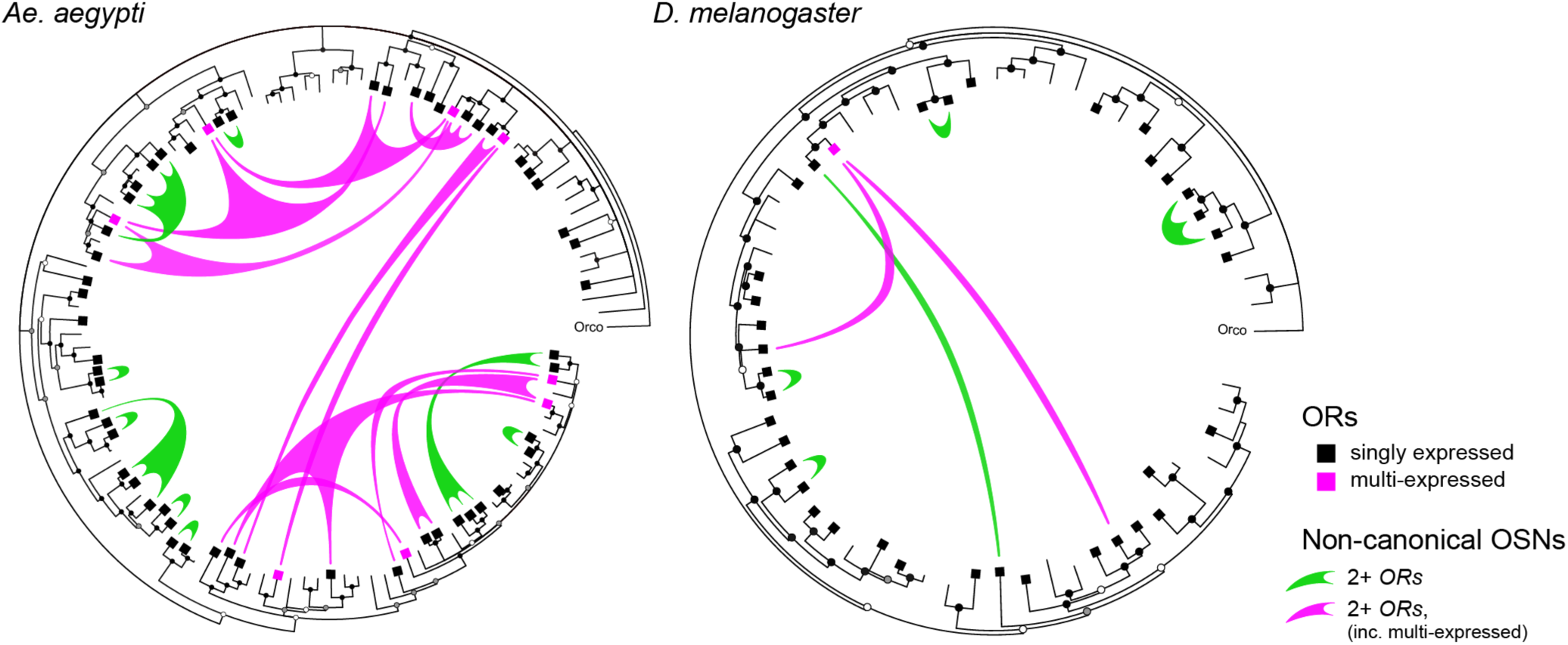
Patterns of *OR* coexpression in *Ae. aegypti* and *D. melanogaster* antennae. Inward circle phylogenies show the evolutionary relationships among *ORs* found in the genomes of the dengue mosquito (left) and vinegar flies (right). Squares mark genes expressed in antennae. Green and pink ‘sails’ represent non-canonical OSN subtypes (i.e. those that express multiple receptors) and extend fingers to each receptor expressed therein, with colors distinguishing subtypes that do or do not express at least one multi-expressed gene (as in Fig. 4A). Taken together, coexpression is much less common in vinegar flies than in the dengue mosquito (fewer sails in tree on right), but shows the same signatures of coexpression by descent among singly expressed receptors (green sails tend to connect closely related ORs) and coexpression by co-option involving multi-expressed receptors (pink sails crisscross the center to connect distant ORs). Trees inferred from protein sequences using the program BAli-Phy (see Methods). Circles at tree nodes indicate bootstrap support: open >0.5, grey >0.7, black >0.9. Receptor expression data for *Drosophila* antennae taken from^3,44,45^.

## Acknowledgements

We thank Richard Benton for comments on the manuscript, Hongjie Li for sharing the antennal nucleus dissociation protocol pre-publication, Tina DeCoste of the Princeton Flow Cytometry Core Facility for help optimizing our FACS protocol, Wei Wang and Jennifer Miller of the Princeton Genomics Core Facility for help with 10X library prep and sequencing, and Vanessa Ruta and members of the McBride laboratory for discussion.

## Author contributions

Conceptualization, EDA and CSM; Investigation, EDA and SK (snRNAseq), EDA, HCM, DT, ZZ and JLZ (CRISPR transgenesis and immunostaining); Data Analysis, EDA, VDA, NHR, and CSM; Writing, EDA, VDA, JLZ, NHR, CSM.

## Materials, data, and code availability

Plasmids generated in this study have been deposited with AddGene (accession #217614, 217647, 217629). Genetically modified mosquito strains are available upon request. Raw snRNAseq reads are available in the NCBI-SRA (BioProject ID PRJNA1138769). Processed snRNAseq data (pre-clustered R objects) and an updated AaegL5 gtf annotation file are available through Zenodo (doi.org<u>/10.5281/zenodo.12797292</u>, doi.org<u>/10.5281/zenodo.12801833</u>). Scripts used for genome reannotation and snRNAseq data analysis are available on github (<u>github.com/mcbridelab/Adavi_2024_snRNAseqAaegAntennae</u>, copy archived at <u>swh:1:rev:ff813a7183ae56cbf3a03a5458450fdf057e674e</u>).

## MATERIALS and METHODS

### Ethics and regulatory information

Human-blood feeding conducted for mosquito colony maintenance did not meet the definition of human subjects research, as determined by the Princeton University IRB (Non Human-Subjects Research Determination #6870).

### Mosquito rearing and colony maintenance

All female mosquitoes dissected in this study were reared at 26°C, 75% RH on a 14:10 light/dark cycle. Eggs were hatched in a broth containing deoxygenated water and Tetramin Tropical Tablets (Pet Mountain, 16110M) and larvae were fed additional tablets *ad libitum* through pupation. Male and female pupae were transferred to plastic buckets or bugdorm cages. Adults were allowed unlimited access to 10% sucrose solution. Where applicable, females were allowed to bloodfeed on a human arm and lay eggs on wet filter paper (Whatman, 09-805B). All transgenic strains used in this study were derived from the Orlando (ORL) laboratory strain except for 15XQUAS-mCD8:GFP strain^33^, which has a Liverpool (LVP) genetic background.

### Generation of *15XQUAS-unc84:2XeGFP* transgenic strain

pBac-mediated transposition was used according to previously published methods^46^. Briefly, the coding sequence of the chimeric protein unc84:2xGFP^17^ was isolated from the original pMUH_unc84_2XGFP plasmid (Addgene #46023) via PCR using the following primers (5’-AACAGATCTGCGGCGGCAAAATGGCTCCCGCAACGGAAG-3’ and 5’-CGGCCCCTAGGGCGGTCACACCACAGAAGTAAGG-3’). The purified insert was then cloned into the AsiSI (NEBioLabs calatolog number R0630) restriction site of the pBac vector (a gift from Leslie Vosshall) which contains the 15XQUAS promoter sequence from the QF2 expression system^47^. Transgene plasmid (500 ng/uL; PBac-15XQUAS-unc84:2XeGFP-3XP3-ECFP, AddGene #217614) was sent to the Insect Transformation Facility, Rockville, Maryland, for microinjection into Orlando (ORL) strain embryos alongside pBac transposase mRNA (300 ng/μL). Out of 256 injected G0 eggs, we recovered over twenty G1 families with 3XP3-ECFP expression in the larval eye, and most remained positive at G2. We outcrossed G2 females to ORL males for several more generations and mapped the number and locations of insertions in each line via the TagMap method^48^. We chose one family (P2) for further use with a single insertion on chromosome 1 far from any annotated coding sequence (Chr1: 61,691,142).

### Generation of Or4-T2A-QF2 and Or103-T2A-QF2 strains

We used CRISPR-mediated homologous recombination^46^ to insert the QF2 transcription factor^47^ into the endogenous *Or4* (AAEL015147) and *Or103* (AAEL017505) loci of the Orlando (ORL) strain of *Ae. aegypti* (Fig. S16). These inserts disrupt the native coding sequences and are thus putative knock-out alleles. We designed sgRNAs targeting the second exon of each locus (*Or4*: GGTGGAGATGATCTACGGTC<u>GGG</u>, *Or103*: GTGCACCTCACGCGCTAG<u>CGG</u>, PAM sequences underlined) and generated dsDNA template for transcription of the sgRNA via template-free PCR with partially overlapping PAGE-purified primers using the NEBNext High-Fidelity polymerase (NEB, M0541S). We then transcribed sgRNA *in vitro* using the HiScribe T7 Kit (NEB, E2040S) with an 8-9 hr incubation at 37°C. We purified the transcription products using RNAse-free SPRI beads (Agencourt RNAclean XP, Beckman-Coulter A63987) and eluted in nuclease-free water.

We constructed the donor plasmid for the *Or4* locus (Or4-T2A-QF2-3XP3-dsRed, AddGene #217647) based on a preexisting plasmid designed for insertion of a different construct at the same locus and cutsite (Or4-T2A-mCD8:GFP-3XP3-dsRed, AddGene #219789). We first amplified the plasmid backbone, including 818bp and 936 bp homology arms and the 3XP3-dsRed marker, but excluding the GSG-T2A-mCD8:GFP element. We then amplified the T2A-QF2-attP construct (1488bp) from different plasmid via PCR, gel extracted and purified both pieces, and cloned them together via InFusion HD cloning (Clontech, 638910). The completed donor plasmid was verified by Sanger sequencing (Genewiz).

We constructed the donor plasmid for the *Or103* locus (Or103-T2A-QF2-3XP3-dsRed, AddGene #217629) using the InFusion HD Kit (Clontech, 638910). Homology arms (∼1 kb each) flanking the Cas9 cut site were amplified from ORL-strain genomic DNA via PCR and cloned into a T2A-QF2-3XP3-dsRed plasmid backbone (linearized with restriction enzymes NsiI-HF (New England Biolabs #R3127S) and AvrII (New England Biolabs #R0174S). The completed donor plasmid was verified by Sanger sequencing.

For each construct, 1500-2000 embryos were injected with a mixture of donor plasmid (700 ng/μL), sgRNA (110 ng/μL) and Cas9 protein (300 ng/μL; PNA Bio, CP01-200) at the Insect Transformation Facility, Rockville, Maryland. A stable transgenic line was developed from one of 19 g1 families (Or4) and from one of 5 g1 families (Or103) showing 3XP3-dsRed expression in the larval eyes. Proper integration into the genome was verified by PCR and Sanger sequencing. Transformed animals were outcrossed to ORL for 5-7 generations, and then maintained by incrossing with continued selection for 3XP3-dsRed+ individuals. Both strains are homozygous viable and show no obvious fitness deficits.

Or4 primers

sgRNA template: forward 5’-GAAATTAATACGACTCACTATAGGTGGAGATGATCTACGGTCG TTTTAGAGCTAGAAATAGC-3’, reverse5’-AAAAGCACCGACTCGGTGCCACTTTTTCAAGTT GATAACGGACTAGCCTTATTTTAACTTGCTATTTCTAGCTCTAAAAC-3’

Amplification of original homology arms from genomic DNA: Left arm forward 5’- CTGTAACTTTCCAGAACACCACATA-3’, Left arm reverse 5’-CGTAGATCATCTCCACCACC-3’, Right arm forward 5’-GTCGGGTGGTTACCGGAG-3’,

Right arm reverse 5’-CGTTTTGTGCGGCAGGTAATAGAG-3’

Amplification of backbone (including homologous arms and 3XP3-dsRed): forward 5’- CGTAGATCATCTCCACCACC-3’, reverse 5’- GATACGCGTACGGCAATTCG-3’

Amplification of T2A-QF2-attP: forward 5’- tggagatgatctacgTGCATGGATCGGGAGAGGG-3’, reverse 5’- tgccgtacgcgtatcGCCGTACGCGTATCTAGAG-3’

Verification of genomic integration: Right arm forward 5’-TGAAGGGCGAGATCCACAAGGC-3’, Right arm reverse 5’-ATGGGCCAAAACTTCCACGCC-3’

Or103 primers

sgRNA template: forward 5’-GAAATTAATACGACTCACTATAGTGCACCTCACGCGCTAGCG GTTTTAGAGCTAGAAATAGC-3’, reverse 5’-AAAAGCACCGACTCGGTGCCACTTTTTCAAG TTGATAACGGACTAGCCTTATTTTAACTTGCTATTTCTAGCTCTAAAAC-3’

Amplification of homology arms: Left arm forward 5’- CAGGCGGCCGCCATATCCCCTTCAAATAGGTAACAATGTATCA-3’, Left arm reverse 5’- CCCTCTCCCGATCCACCCAGGGCGTGCGGGGCCGAGAAAGCTATTTCAGTTGCCTTATTCGGGATT-3’; Right arm forward 5’- TGTATCTTATCCTAGGCGTGGACTAATAAATATGGATCAGCAATT-3’, Right arm reverse 5’-TATTAATAGGCCTAGGGACTTATGAGACTTATATTGATCATGTACTTCTCA-3’.

Verification of genomic integration: Left arm forward 5’-GCCCAAACCCGTACGGTAATAA-3’, Left arm reverse 5’- CGTAGTTGTGGGTCCCAGAC-3’, Right arm forward 5’- CGGCCGCGACTCTAGATCATAATCAG-3’, Right arm reverse 5’- GCAAAGCTATACTAAAATAAAACATCGGGACT-3’

### Visualization of Or4 and Or103 target glomeruli via brain immunostaining

*Or4-T2A-QF2* or *Or103-T2A-QF2* animals were crossed with *15XQUAS-mCD8:GFP* animals (Liverpool background^47^), and offspring were screened for those inheriting both constructs. Brain immunostaining was carried out as previously described^47^ on 7–10 day-old mated female mosquitoes. Heads were fixed in 4% paraformaldehyde (Electron Microscopy Sciences, 15713-S) for 3 hours at 4°C. Brains were dissected in PBS and blocked in normal goat serum (2%, Fisher Scientific, 005-000-121) for 2 days at 4°C. We then incubated brains in primary antibody solution for 2–3 days, followed by secondary antibody solution for another 2–3 days at 4°C. Brains were mounted in Vectashield (Vector, H-1000) with the anterior side facing the objective. Confocal stacks were taken with a 20X lens with XY resolution of 1024X1024 and Z-step size of 1 mm. Primary antibodies: rabbit anti-GFP (1:10,000 dilution, ThermoFisher, A-11122) and mouse NC82 (1:50 dilution, DHSB, AB_2314866). Secondary antibodies: goat-anti-rabbit Alexa 488 (1:500 dilution, ThermoFisher, A27034SAMPLE), goat-anti-mouse CF680 (1:500 dilution, Biotium, 20065-1).

### Female antennal dissection

We crossed the *brp-T2A-QF2w*^11^ driver to our new *15XQUAS-unc84:2XeGFP* effector and screened offspring as larvae for those inheriting both constructs (3XP3-dsRed and 3XP3-ECFP positive). Males and females were co-housed with access to 10% sucrose but not blood. Female mosquitoes were dissected at 5–7 days post eclosion. To ensure a sufficient nuclei yield for downstream sorting, we designed a dissection protocol to remove and preserve the integrity of the antennal flagellum (excluding the pedicel). First, mosquitoes were anesthetized at 4°C for 25 minutes and 70 individuals were placed on a prechilled Sylgard® coated 93 mm diameter petri dishes (Living systems instrumentation, DD-90-S). The dissection dish was placed on ice and transferred to a stereoscope at room temperature. Under the stereoscope, we severed the head of each female one at a time with sharp forceps, swirled it gently in a dish of pure ethanol for 3-5 seconds at room temperature, rinsed it twice with vigorous swirls in large amounts of room temperature Dulbecco’s PBS without calcium chloride and magnesium chloride (Sigma-Aldrich, D8537), and then placed it neck down on a second prechilled Sylgard® coated petri dish on ice containing Schneider’s *Drosophila* Medium (Thermo Fisher, 21720024). When working well, cuticular wax can be seen diffusing away from the head during the dip in ethanol. Once all heads were collected, we placed the second Sylgard coated petri dish on ice under the stereoscope to remove both antennae from each head by (1) holding the head with one forceps, (2) grabbing the base of the flagellum (first flagellomere) with the other forceps, (3) pulling it away from the head, and (4) releasing it in the Schneider’s medium within the petri. With sharp enough forceps, the antennal pedicel, which contains the Johnston’s organ, stays attached to the head, and we confirmed that all dissected antennae were free of pedicel residue. We then used a pipet with wide orifice pipette tips (Thomas Scientific, 1234W) to transfer the flagella from the petri into a 1.5mL non-stick microcentrifuge tube (Neta Scientific, RPI-145530) on ice (Thomas Scientific, 1234W). The antennae naturally sink to the bottom of the tube after 5-7 minutes, after which the flagella can be concentrated by pipetting off the liquid. This procedure was performed simultaneously by two people and both pools of dissected flagella were merged into a single microcentrifuge tube to double the yield of tissue per tube/session. Finally, the flagella were instantly snap frozen in a liquid nitrogen-cooled mini mortar (SP Bel-Art, H37260-0100) and stored at -80°C until nuclear isolation. The entire procedure lasted 45-50 minutes and was repeated approximately 30 times on different days with different batches of mosquitos.

### Nuclear isolation

We followed a previously described method for isolating *Drosophila* nuclei^49^ with minor modifications as follows: two microcentrifuge tubes of frozen antennae were transferred to the bench in a -20°C cooling block (one at a time), 100 µl of prechilled HB lysis buffer (250mM Sucrose, 10mM Tris pH 8, 25mM KCl, 5mM MgCl2, 0.1% Triton-x 100, 0.5% RNAse inhibitor Plus (Promega, N2615), 1x Protease inhibitor cocktail dissolved in DMSO (Promega, G6521), 0.1mM DTT) were quickly added to the tube outside of the cooling block and antennae were immediately pulverized with a motor pestle for 30s next supplemented with 400µl HB. Both batches were merged into a single glassTissue Grinder (Wheaton®, 357538) for a total volume of 1mL. The suspension was homogenized with 18 loose Dounce strokes followed by 26 tight Dounce strokes on ice. The lysate was strained through a 35µm meshed 5mL test tube (Corning® Falcon®, 35223535) to remove cuticular aggregates, filtered through a prewet 40µm Flowmi™ cell strainer (Bel-Art, H13680-0040) to remove cellular clumps, and centrifuged at 1000g in swinging buckets (Eppendorf, Rotor S-24-11-AT) at 4°C for 10 minutes.

After centrifugation, as much supernatant as possible was removed from the tube and the nuclear pellet (often visible in shades of white to dark grey) was resuspended by pipetting up and down 20 times in 400µL of Washing Buffer (1X Dulbecco’s PBS without calcium chloride and magnesium chloride, 1% MACS BSA stock solution (Miltenyi Biotech,130-091-376), 0.5% RNAse inhibitor Plus). During the centrifugation time, the procedure was repeated on a second round of two batches. During the centrifugation of the second round, the first round was resuspended and then kept on ice until the resuspension of the second round. Once both were resuspended, each isolate was individually filtered through a prewet 40µm Flowmi™ to remove nuclear clumps and collected into the same 1.5mL non-stick microcentrifuge tube (Neta Scientific, RPI-145530) for a total volume of 750-800µL depending on Flowmi™ retention. Hoechst33342 (Thermo Fisher Scientific, 62249) was added 5–10 minutes before sorting at 1µg/ml final concentration. The suspension was kept on ice at all times until FACS loading.

### FACS sorting & nuclear imaging

The full 750-800µL suspension was loaded at 4°C into a FACSAria™ Fusion flow cytometer (BD Biosciences) with a 70 µm nostril running regular 1X PBS at the Flow Cytometry Core Facility of the Molecular Biology Department at Princeton. We empirically refined the optical parameters to isolate healthy single nuclei as follows. We used the first gate [FSC-A(exponential) X DAPI-A(linear x1000)] to select all events in the first of two Hoechst intensity peaks. Most nuclei in the second peak were confirmed by imaging to correspond to doublets or single nuclei with unusually bright Hoechst intensity (possibly in G2/S phase, data not shown). We used a second gate [SSC-W(linear x1000) X SSC-H(exponential)] to discard additional potential doublets or aggregates. We used a third gate [SSC-A(exponential) X GFP FITC-A(exponential)] to select and separate GFP positive neuron nuclei (∼35%) from GFP negative nuclei (∼65%). We used a fourth gate [DAPI-A(linear) X SSC-A(exponential)] to remove GFP nuclei with heterogeneous granularity which corresponded to damaged single nuclei. This procedure was conducted on three different nuclear suspensions, each processed on a different day and composed by different antennal dissection batches. Together, the three FACS runs yielded 7 samples of approximately 20,000 GFP positive nuclei each (1 sample from the first run and 3 samples from each of the second and third runs). The samples were collected in nonstick 1.5ml microcentrifuge tubes smeared with 12 μl fresh prechilled Washing Buffer.

### Single nuclei RNA sequencing

We measured the volume of each sample, added up to 43µl of nuclease-free water, and loaded it into a 10X Genomics Chromium system using Reagent Kits to generate and amplify cDNAs as recommended by the manufacturer (10X Genomics). We checked the electrophoretic profile of cDNA libraries before and after tagmentation, selected fragments of size 300-700bp using a BluePippin (Sage Science), and generated 100 bp paired-end sequencing reads on an Illumina NovaSeq 6000 SP flowcell following standard Illumina protocols. Only pass-filter reads were retained by Illumina Control Software and were aligned to the *Aedes aegypti* L5 genome excluding extrachromosomal nuclear contigs^50^ with CellRanger v6.0.1 with the “include-introns” option on (10X Genomics).

### Genome Reannotation

We made several custom updates to the publicly available AaegL5 genome annotation (VectorBase-55_AaegyptiLVP_AGWG.gff) to facilitate analysis and interpretation of our data. First, we removed the few annotations present for small ‘unplaced’ contigs, leaving only annotations for the three major chromosomal scaffolds and one mitochondrial scaffold. Second, since the annotation was missing several chemoreceptors that might be expressed in female antennae, we revised it to include the manually curated chemoreceptor (OR, GR, IR) annotations provided in the supplement of Matthews et al. 2018^50^. This involved (i) renaming the contigs from the manual annotation file to match those use by VectorBase, (ii) identifying genes in the main gff file that overlapped those in the manual chemoreceptor gff file using sed and bedtools intersect (-wa -u), (iii) removing from the main file all features associated with these genes by feeding grep (-Fvf) a list of gene_ids, (iv) merging the two gff files, and (v) converting to gtf using gffread (-T -o)^51^. Finally, we used metazoa.ensemble.org^52^ to infer potential *Drosophila* orthologs for as many mosquito genes as possible and appended the *Drosophila* names to the ends of the mosquito names where possible.

We also developed an automated pipeline that leveraged our snRNAseq data to extend 3’ UTRs in the AaegL5 annotation (Fig. S2). Incomplete annotation of 3’ UTRs is a widely recognized problem for the analysis of 3’ sequencing data such as that generated here using the 10X genomics platform^53,54^.

Reads that pile up just downstream of an annotated UTR will not be assigned to the proper gene, causing loss of signal. Briefly, we first extracted the subset of 10X reads that aligned to small windows starting 100bp upstream of the 3’ end of any annotated transcript or transcript-like feature and extending 750bp downstream. We then used StringTie (-p 40 -m 30 --j 1000000000 --fr)^55^ to assemble these reads into short ‘gene models’, with each ‘gene model’ representing a pile-up of 10X reads just downstream of an annotated gene. Finally, we used a custom python/gffutils script to extend UTRs to the end of any StringTie ‘gene model’ present within its +750bp window. If the extended UTR overlapped with a downstream neighbor on the same strand, we truncated it immediately upstream of that neighboring gene. We used gff3sort.pl^56^ to sort the updated annotation and gffread (-E -t -o) to again convert to gtf format.

We visually inspected read alignments for ORs and IRs and manually extended them up to 5 kb when the read trace continued further downstream of the 750bp window and/or included one or more extra peaks. For each extra peak, we looked for alternative polyadenylation sequences (AAUAAA) and extended the 3’UTR to either the first A of the alternative sequence (when present) or the last nucleotide of the last mapping reads of the last extra peak.

The final updated gtf file is available at https://doi.org/10.5281/zenodo.12801833.

### Data preprocessing, ambient RNA decontamination and doublet removal

CellRanger-generated UMI count matrices were loaded into singleomics R toolkit Seurat 4.2.0^57^. We merged all 7 libraries and discarded all droplets with low complexity (less than 350 genes detected), high UMI counts (>7000 UMIs detected), and/or high mtDNA content (>0.5% reads). We normalized all droplets together using the Seurat command SCTransform with v2 regularization (vst.flavor=”v2”)^58,59^, ncells=47388 (equal to the total amount of droplets in the dataset to avoid droplet subsampling), n_genes=10000 (up from 2000 to reduce gene subsampling) and all others at default values. We then explored a variety of clustering parameters in order to identify the most robust. We varied the number of PCs (npcs in the RunPCA) between 10 and 200. We varied the number of dimensions (dims in the RunUMAP) to match the number of PCs in a given run. We varied the number of neighbors (n.neighbors) between 10 to 200. For the FindClusters command, we set the resolution to 1 and chose SLM (algorithm=3) instead of the default Louvain algorithm. Visual comparison of the resulting UMAPs revealed that the number and identity of clusters (i.e. putative cell types) was particularly sensitive to the number of PCs. More specifically, the number of clusters increased non-linearly with the number of PCs up to a saturation point after which the number of clusters remained stable. We therefore chose to use the lowest number of PCs that allowed us to reach the saturation point and then the number of neighbors that provided the best visual segregation pattern given that number of PCs. The final normalization and pre-clustering parameters were as follows: SCTransform(n_cells=47388, n_genes=10000, vst.flavor=“v2”); RunPCA(npcs=60); RunUMAP(reduction=“pca”, dims=1:60, n.neighbors=110); FindNeighbors(reduction=“pca”, dims=1:60); FindClusters(resolution=1, algorithm=3).

We removed ambient RNA contamination using SoupX (autoEstCont maxMarkers=5000)^60^. We ran SoupX on each library individually to account for library-specific RNA contamination. However, we fed the program cluster identities derived from the unified “pre-clustering” analysis of all 7 libraries described above. After rounding the corrected matrices for each library, we saved them using DropletUtils::write10xCounts^61^ and removed all predicted doublets with the Python package Solo^62^ (-p, using recommended model_json parameters). We finally merged all 7 corrected count matrices and obtained a final data matrix with 46073 curated droplets. We further explored the quality of the preprocessing by comparing dispersion, cumulative distribution, and residual variance between pre- and post-processed data.

### All neuron normalization and clustering

We normalized the curated data matrix using the Seurat command SCTransform as described for pre-clustering except we raised the default number of genes to the total number of genes, scaled and centered the residuals, and reduced maximum residual variance to 50, which was empirically defined. We found that reducing the clipping range was essential to avoid artifactual clusters caused by specific genes with unusually high residual variance (e.g. *nompC*, Fig. S4D). The final normalization parameters were as follows: ncells=total_cells, n_genes=NULL, vst.flavor=“v2”, do.scale=TRUE, do.center=TRUE, clip.range=(-50,50), variable.features.rv.th=1.3. We then clustered the data, again as described for pre-clustering except that we ran UMAP with the uwot method and cosine metric. The final clustering parameters were as follows: RunPCA(npcs=60); RunUMAP(reduction=“pca”, dims=1:60, n.neighbors=110, umap.method=”uwot”, metric=”cosine”); FindNeighbors(reduction=“pca”, dims=1:60); FindClusters(resolution=1, algorithm=3).

### THSN and OSN renormalization and clustering

After generating the “all neuron clustering”, we excluded four ‘junk’ clusters (low complexity clusters in the middle of the UMAP; Fig. 1D #1, 47, 48, 55), two *nompC+* mechanosensory neuron clusters (#29, 54), and one unidentified cluster (#45) before subsetting the remaining clusters into three categories based on *orco* and *Ir93a* expression: thermo/hygrosensory neurons (THSNs; *Ir93a+*, *orco-*), *orco+* olfactory sensory neurons (*orco+* OSNs; *orco+*, *Ir93a-*), and *orco-* olfactory sensory neurons (*orco-* OSNs; *orco-*, *Ir93a-*). We then independently renormalized and reclustered the nuclei belonging to each of the three types of clusters with different parameters based on QC and a clustering sensitivity analysis similar to that described for *Data Preprocessing* (above). Notably, we used 12, 20, and 60 PCs when running RunPCA on THSNs, *orco-* OSNs, and *orco+* OSNs, respectively, to reflect the differing levels of complexity of these groups of cells. We also reduced the number of neighbors for THSNs and *orco-* OSNs to 15 and for *orco+* OSNs cluster (n=59 droplets) represented a mix of three other *orco+* OSN clusters. Importantly, it ‘co-expressed’ groups of ORs that did NOT show correlated expression at the droplet level. We therefore removed these droplets and renormalized/reclustered the remaining *orco+* OSN data a second time. We conducted a similar cleaning step for *orco-* OSN clusters by removing one cluster with putative contamination from orco+ cells (n=248 droplets) and and another very small cluster (n=85 droplets). After renormalization, we again removed the smallest cluster (n=61 droplets), which was a small offshoot of the *Amt+* cluster likely caused by a statistical artifact.

Inspection of the UMAPs and receptor expression patterns revealed heterogeneity within a few OSN clusters. In these cases, we looked for evidence of multiple underlying neuron subtypes using the FindSubCluster command. This resulted in the splitting of 1 *orco+* OSN cluster (#21) and 2 *orco-* OSN clusters (#10 and 14). See Fig. S9 for visualization of how the original ‘all neuron’ clusters relate to those in the reanalyzed *orco+* OSN and *orco-* OSN subsets.

### Quantification of co-receptor expression in all neuron clustering

We examined the distribution of average expression values for four olfactory co-receptors (*orco*, *Ir25a*, *Ir8a*, *Ir76b*) across clusters in the all-neuron analysis (Fig. 1E) and identified a natural break as the on-off cutoff: each co-receptor was considered expressed if its average expression in a given cluster exceeded 10% of the gene-specific maximum across all clusters.

### Quantification of receptor expression in THSN, *orco+* OSN, and *orco-* OSN clusters

We quantified receptor expression within clusters using a median-adjusted log2 fold change (log2FC’). Log2FC compares expression in a focal cluster to expression in all other clusters. The adjustment was made by subtracting the median log2FC value across all clusters from the log2FC in focal clusters. We made this adjustment because low-level background expression led to many low (but positive) values for some receptors. The distribution of log2FC’ values across all receptor-by-cluster combinations (Fig. S12A, S15C) revealed clear breaks or inflection points that we used as on/off thresholds (0.4 for *orco-* OSNs, 0.3 for *orco+* OSNs). Alternative absolute log average expression thresholds produced similar results (0.15 for *orco-* OSNs, 0.16 for *orco+*; Fig. S12C, S15B).

More than one ligand-specific receptor was called as expressed in many OSN clusters. To confirm co-expression, we looked for correlated expression at the droplet level and found significantly elevated Pearson correlations in all but one case. *Ir41b* was called as expressed in *orco-* OSN cluster 10b, alongside *Ir41e* and *Ir41l*, but showed no sign of correlated expression with the latter two receptors at the droplet level.

In a few cases, two OSN clusters expressed the same set of receptors (e.g. *orco-* OSN clusters 14a/14b, 4/9 and *orco+* OSN clusters 15/37, 21a/21b, 21c/24). These may represent (i) distinct OSN subtypes that express the same set of receptors, (ii) the same OSN subtype from individuals with different genotypes (given the presence of genetic variation within the ORL lab strain) or in different biological states, or (iii) clustering artifacts. Conservatively, we decided to merge such clusters for the purposes of downstream analyses.

### Receptor gene tree inference

We inferred phylogenetic trees for the *Ae. aegypti IR*, *Ae. aegypti OR*, and *D. melanogaster OR* families using Bali-phy 3.6.1^63^. We first tested different parameter combinations and settled on the following substitution frequencies, heterogenous rates and indel models for all gene families: -S wag+f+Rates.gamma+inv -I rs07. We then (1) ran ten simultaneous iterations of 50000 generations each, (2) merged all iterations discarding the minimum burnin, (3) calculated posterior probabilities for all nodes, and (4) only retained phylogenetic relationships with probabilities above 50%. Trees were visualized as inward phylogenies using the Rstudio package ggtree^64^. Phylogenetic distances between pairs of receptors was computed using the cophenetic.phylo function in the Rstudio package phytools^65^.

## SUPPLEMENTAL FIGURES

**Figure S1.**
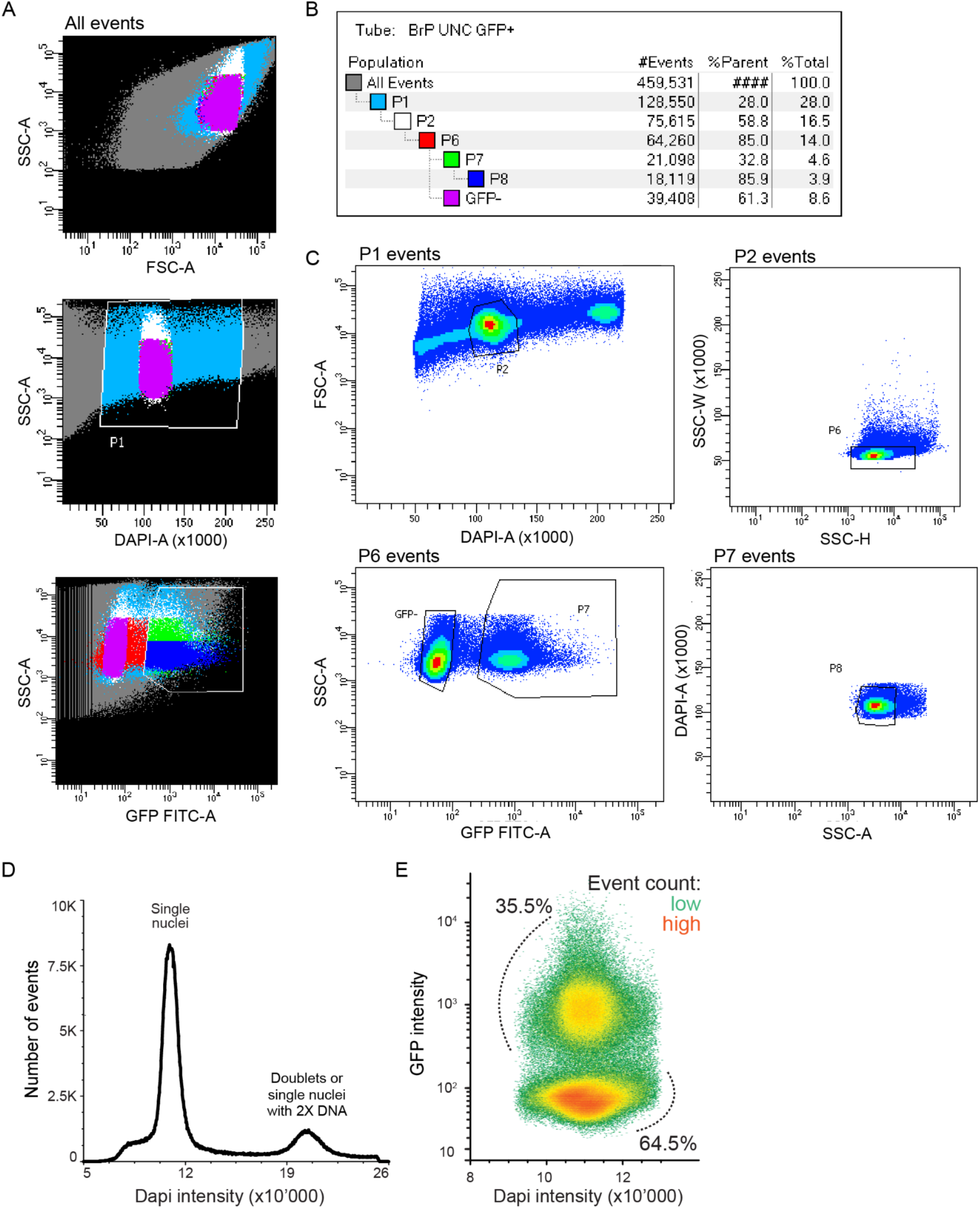
Fluorescent activated cell sorting of GFP-labeled nuclei from female antennal neurons. (**A**) Merged FACS data from 3 independent suspensions of antennal nuclei colored according to the populations shown in (B). (**B**) Table showing the number of events in subsets defined by sequential gates P1-P8 during one FACS run. P8 represents the final subset of GFP+ single nuclei used for 10X library prep. (**C**) Density plots showing sequential gating parameters for one FACS run. (**D**) Distribution of DAPI intensity across all P1 events. Events in the larger 1X peak were retained through gate P2 while those in the smaller 2X peak were discarded. (**E**) Density plot showing all P6 events according to DNA content (DAPI) and GFP intensity. Events in the upper GFP+ peak were retained through gate P7 while those in the lower GFP-peak were discarded. The retained GFP+ neurons represent ∼35% of all antennal nuclei. FSC, forward scatter; SSC, side scatter; GFP-FITC, GFP intensity; DAPI, DNA marker intensity. -A, -W, -H correspond to average, width, height components.

**Figure S2.**
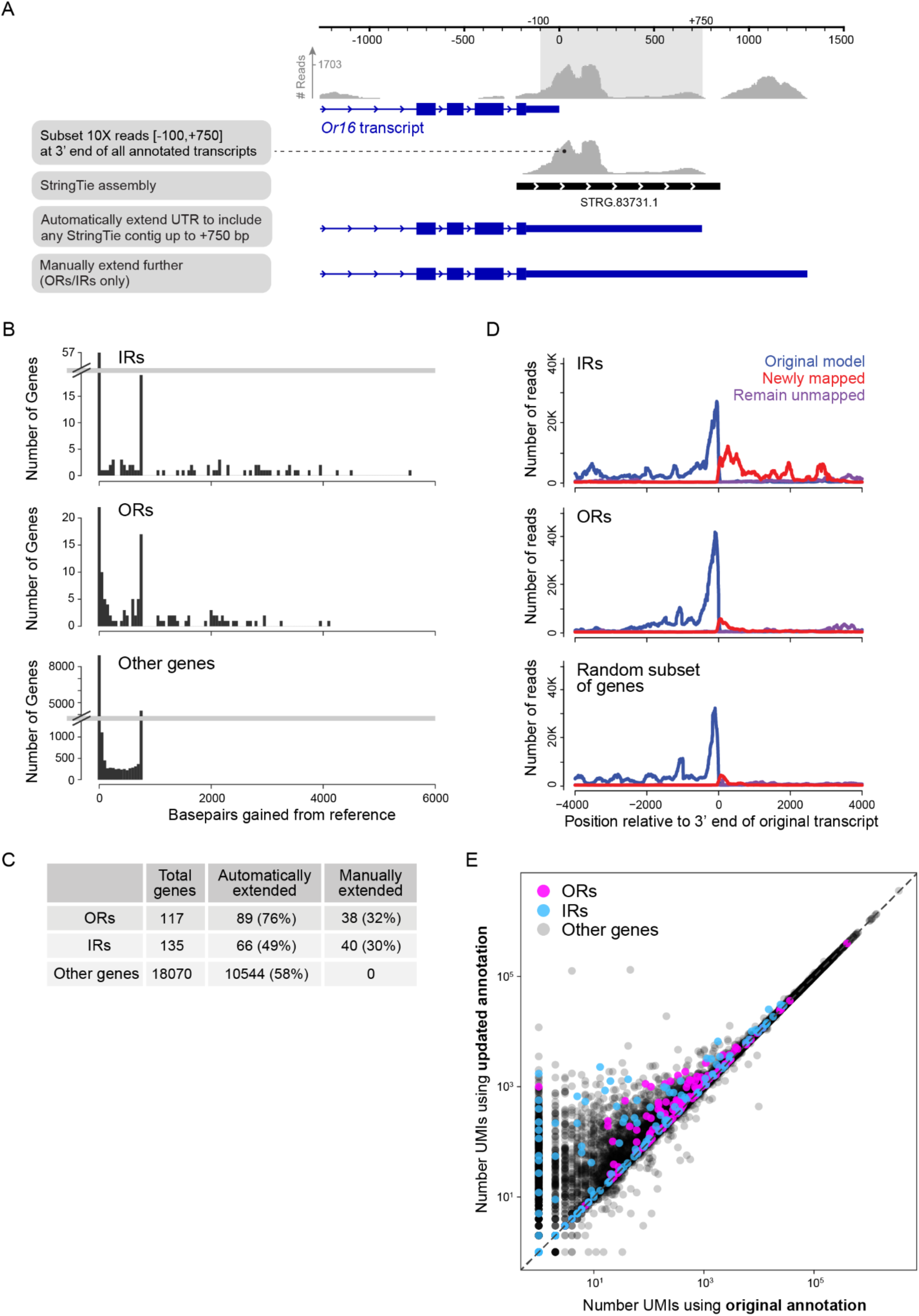
Reannotation or 3’UTRs in the AaegL5 genome. (**A**) Schematic of reannotation process for an example odorant receptor. Top trace shows pile up of 10X reads with base pair coordinates given in reference to the end of the originally annotated transcript. (**B**) Number of base pairs added to the 3’ UTRs of ORs, IRs, and other genes. Many genes in all three categories were extended by exactly 750 bps as this was the extension cutoff used in the automated pipeline. (**C**) Number and fraction of genes that received extensions. (**D**) Distribution of summed 10X reads across the end of annotated ORs, IRs, or a random subset of genes. Colors highlight reads that were assigned to genes using the original annotation (blue), reads that were newly assigned using the updated annotation (red), and reads that were not assigned to the focal genes. (**E**) Scatterplot comparing the total number of UMIs assigned to given genes using the original and updated annotations.

**Figure S3.**
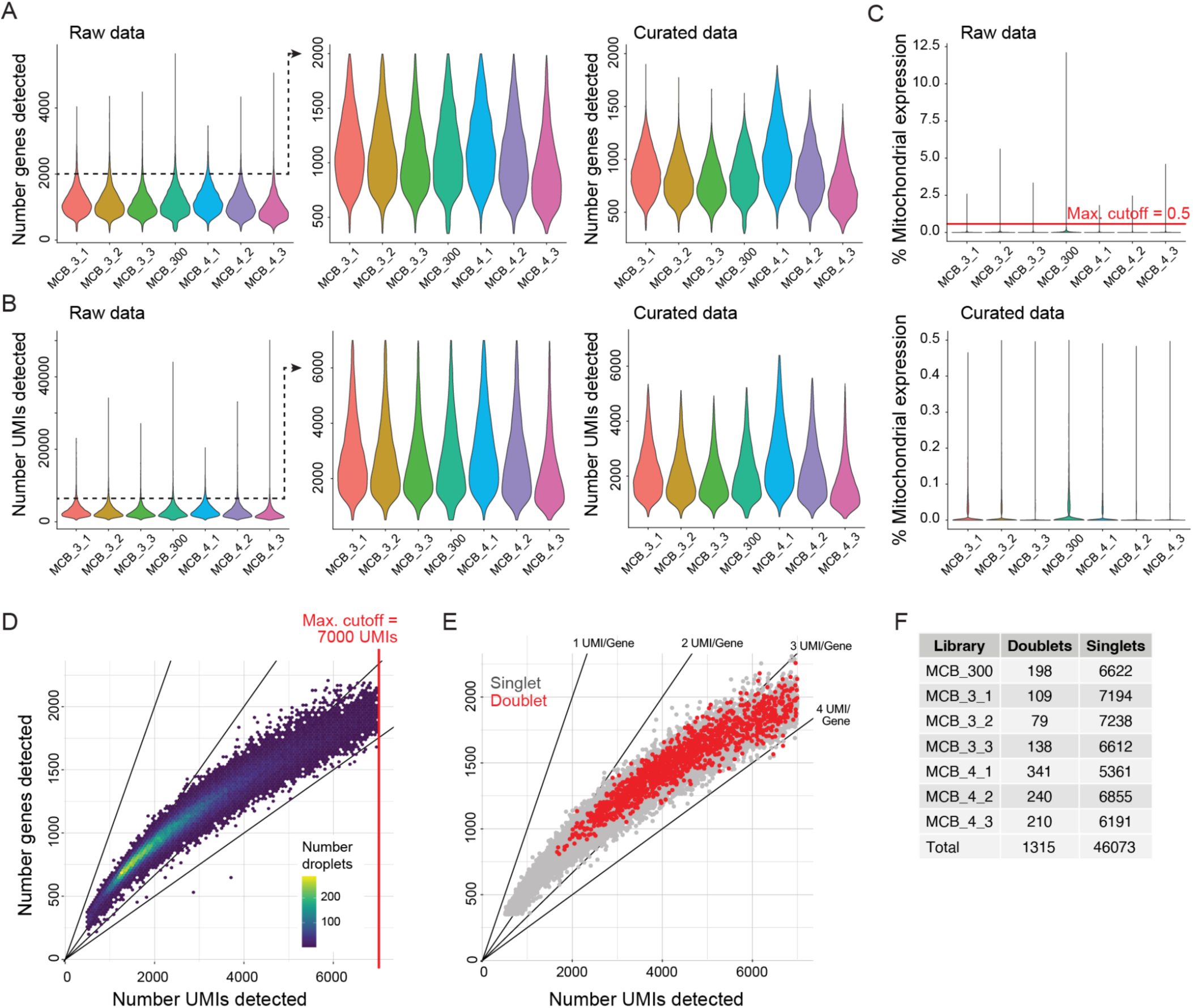
Preprocessing of droplets in 7 replicate snRNAseq libraries. (**A**–**C**) Distribution of genes detected (A), UMIs detected (B), and percent mtDNA transcripts (C) in each library before (raw) and after (curated) removal of ambient RNA contamination using the program SoupX^60^. Droplets with <350 genes, >7000 UMIs, or >0.5% mtDNA expression were discarded. (**D**) Density of droplets in which specific combinations of UMIs and genes were detected. (**E**) As in (D) but highlighting putative doublets detected using the program Solo^62^. (**F**) Summary of doublets detected in each library.

**Figure S4.**
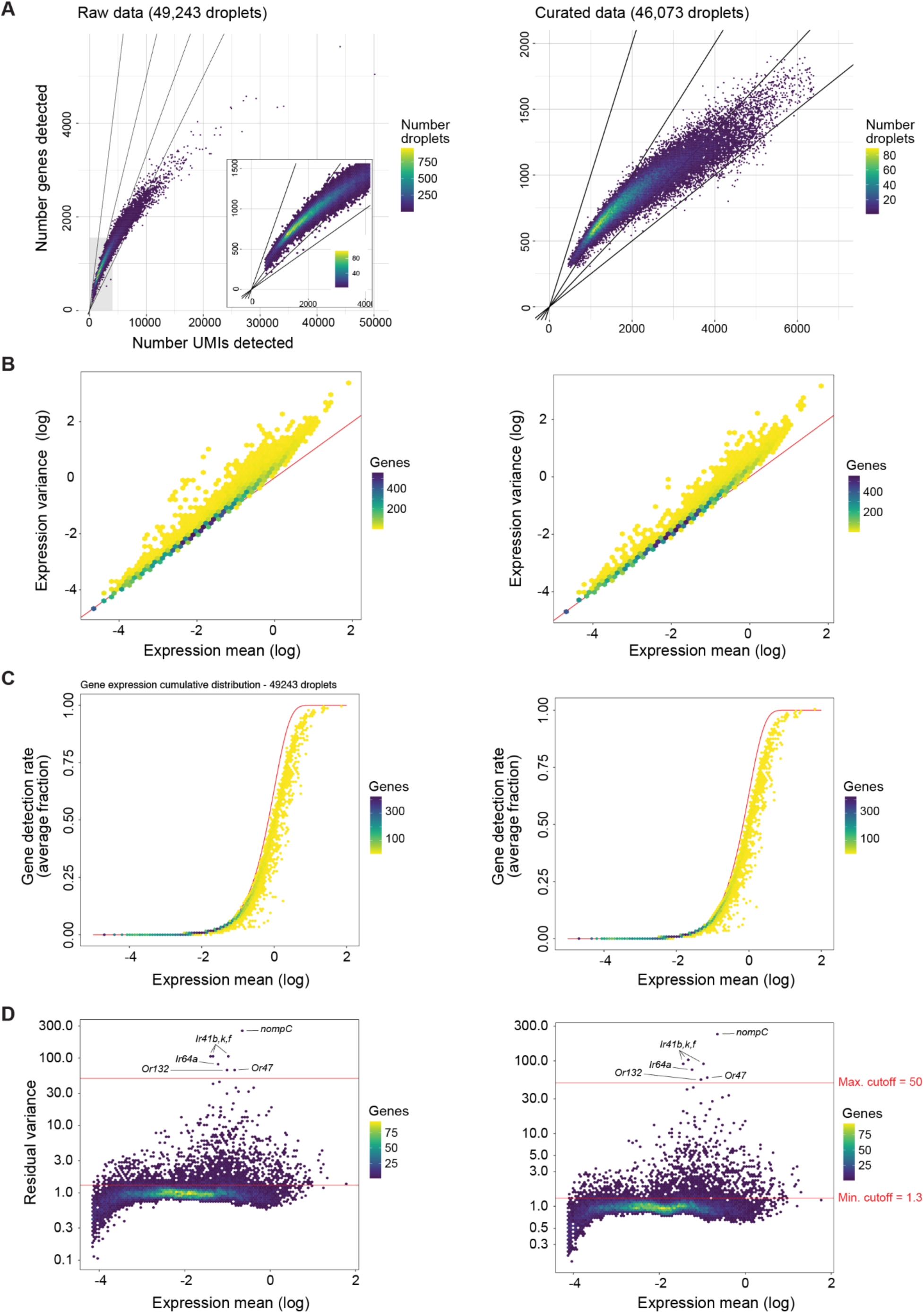
Gene expression variability and normalization in raw and processed droplets from the full dataset. (**A**) Average UMI per gene per droplet. (**B**) Linear modeling of gene expression dispersion. (**C**) Poisson modeling of gene expression deviation. Red line depicts the predicted cumulative distribution. (**D**) Residual expression variance by mean expression across all genes. The highlighted clip.range and rv.th threshold values were used for sctransform v2 normalization. In all panels, plots on the left and right show raw and processed data, respectively.

**Figure S5.**
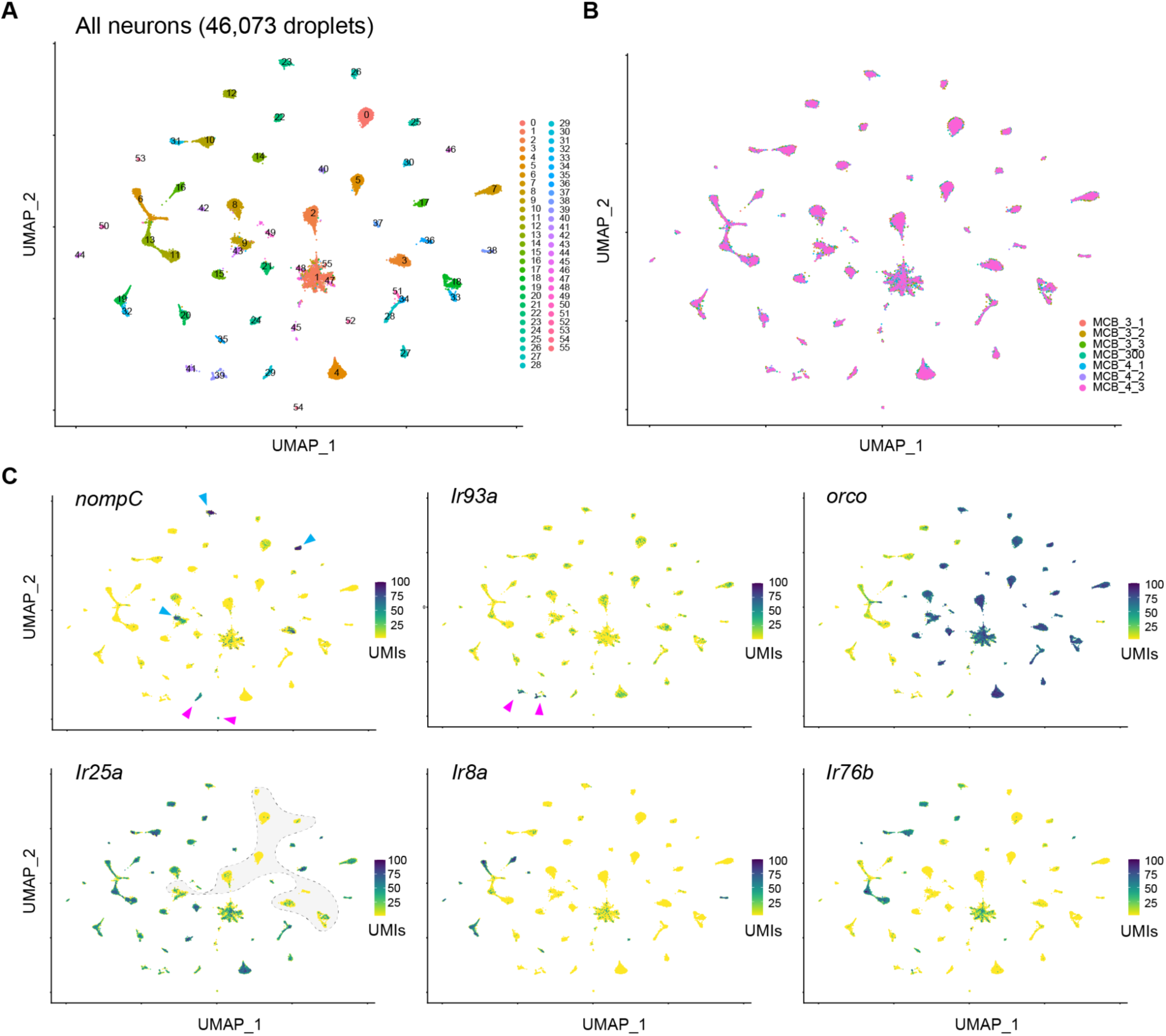
Details of UMAP clustering and marker gene expression in the all-neuron analysis. UMAP clustering for the full data set (as in Fig. 1D) highlighting clusters (**A**), 10X libraries (**B**), and expression of key marker genes (**C**). In (A), clusters 1, 47, 48, and 55 were low complexity ‘junk’ clusters located in the center of the plot, while cluster 45 was clearly distinct but could not be associated with a specific function. In (C), pink arrowheads indicate putative mechanosensory neuron clusters (*nompC* panel), and thermo/hygrosensory neuron clusters (*Ir93a* panel). The mechanosensory channel *nompC* is also highly expressed in several *orco+* olfactory sensory neuron clusters (blue arrowheads). The biological significance of this expression is unknown.

**Figure S6.**
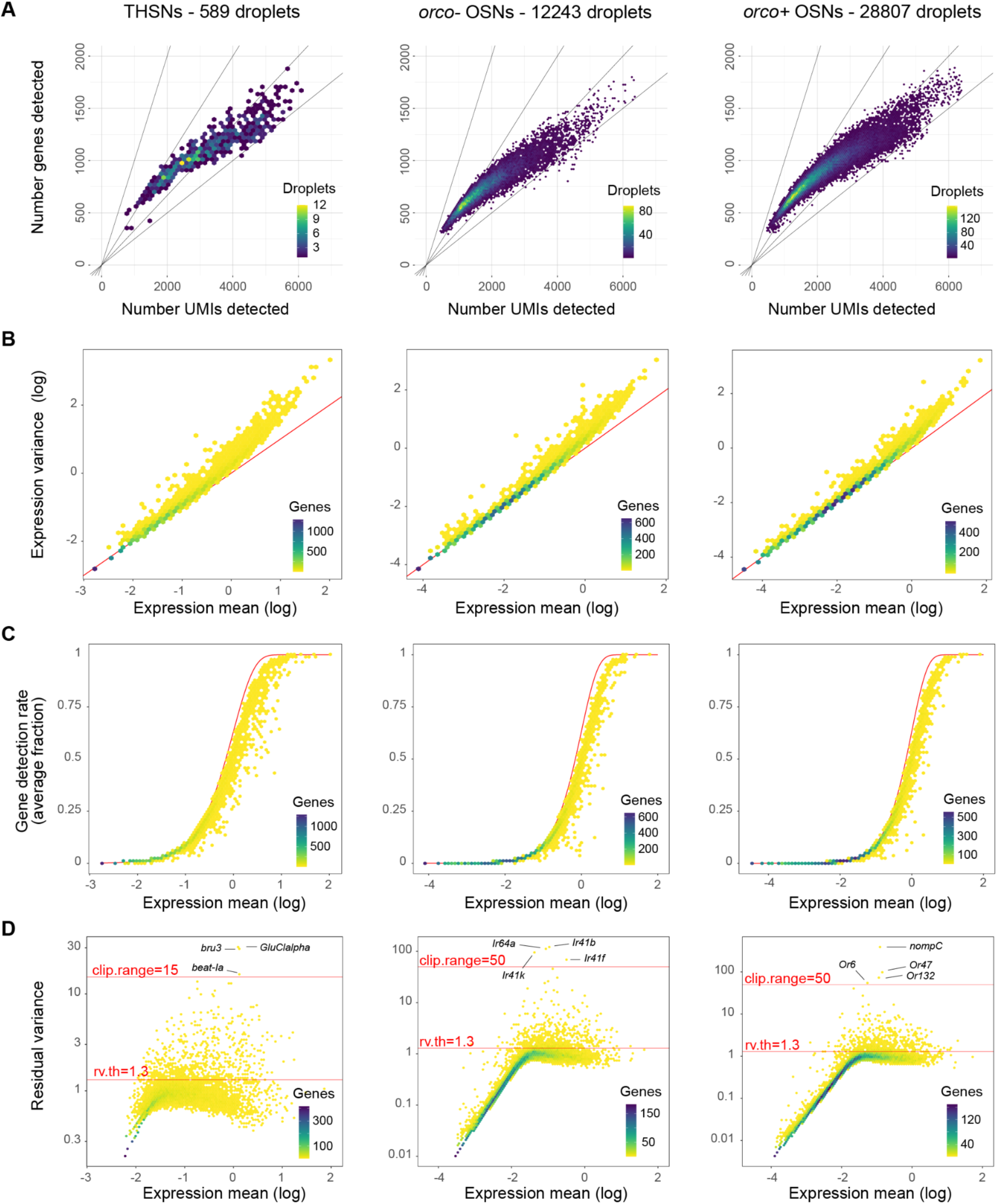
Gene expression variability and normalization of subsetted sensory neurons. (**A**) Average UMI per gene per droplet. (**B**) Linear modeling of gene expression dispersion. (**C**) Poisson modeling of gene expression deviation. Red line depicts the predicted cumulative distribution. (**D**) Residual expression variance by mean expression across all genes. The highlighted clip.range and rv.th threshold values were used for sctransform v2 normalization. In all panels, plots on the left and right show raw and processed data, respectively. Left, middle, and right plots represent thermo/hygrosensory neurons, *orco-* OSNs, and *orco+* OSNs, respectively.

**Figure S7.**
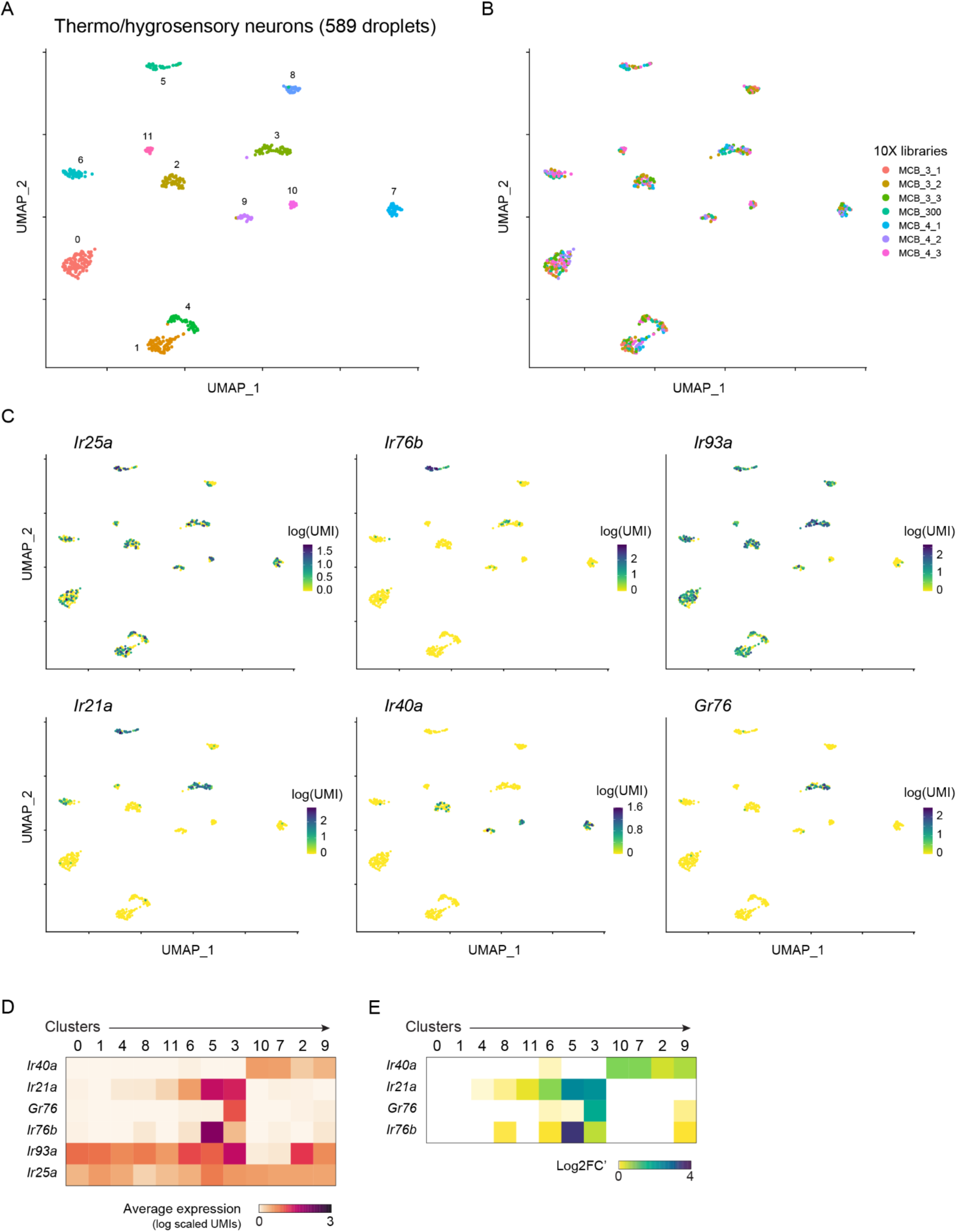
Details of UMAP clustering and receptor expression in putative thermo- and hygrosensory neuron subtypes. (**A**–**C**) UMAPs highlight nuclei assigned to different clusters (A), derived from different 10X libraries (B), or expressing different levels of key chemosensory receptors and coreceptors (C). (**D**–**E**) Heatmaps showing average expression (D) and differential expression (E) of chemosensory receptors across all clusters.

**Figure S8.**
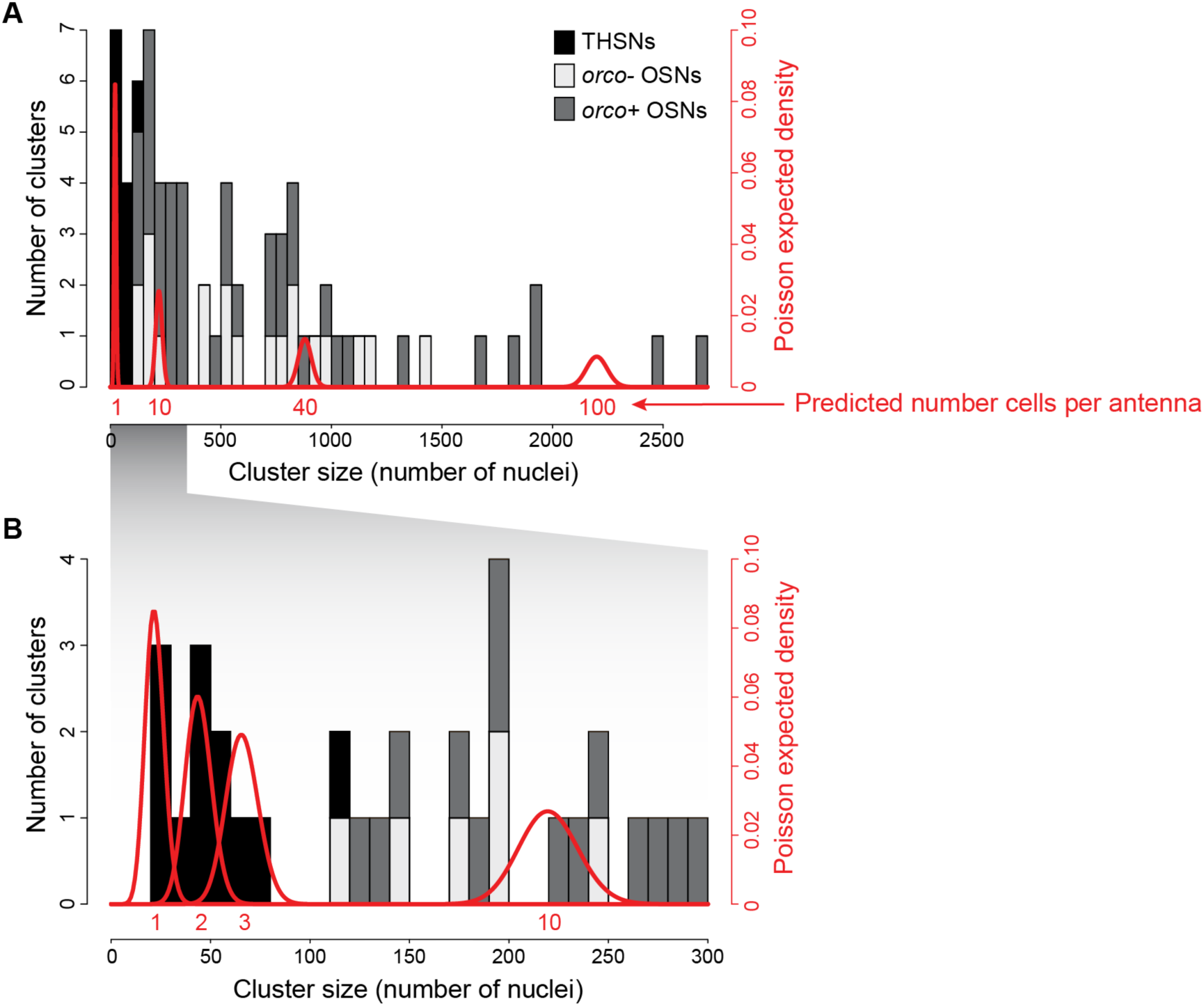
Inferred number of cells on a single female antenna belonging to identified sensory neuron subtypes. (**A**) Histogram showing size distribution of sensory neuron clusters (grey) overlaid by the expected densities for cell types comprising 1, 10, 40, or 100 neurons per antenna (red). Poisson densities were estimated under the assumption that the 46,073 sequenced nuclei were randomly drawn with replacement from the ∼2000 neurons present on a female antenna^19^. (**B**) Zoom of (A), highlighting tiny populations of thermo/hygrosensory neurons (black) that we infer to comprise just 1-3 neurons per antenna.

**Figure S9.**
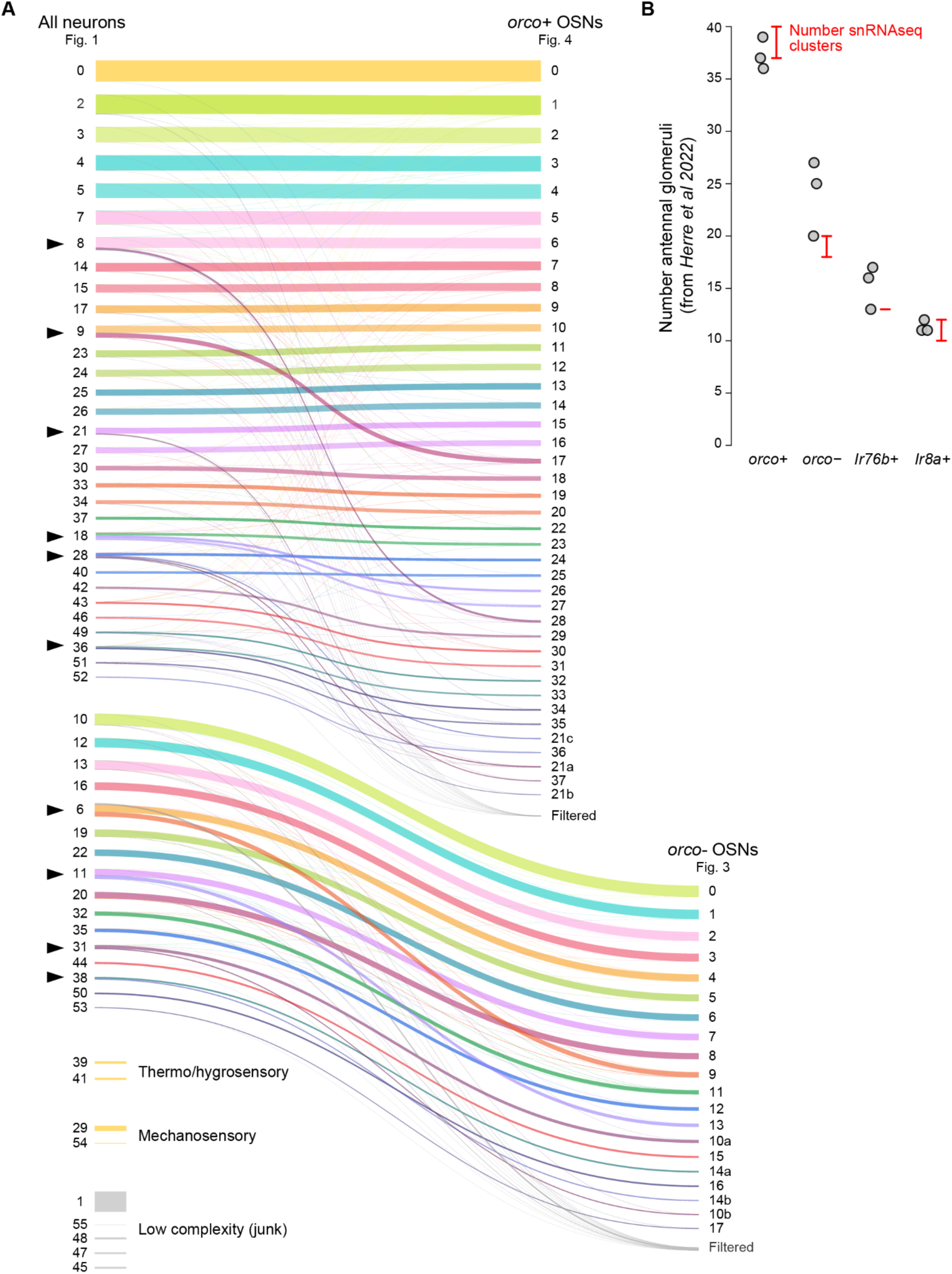
Correspondence between clusters in full and subsetted analyses with comparison to glomerulus counts. (**A**) Sankey plot showing how clusters in the original all neuron analysis (left; Fig. 1D) correspond to those in subsetted *orco+* OSN analysis (top right; Fig. 4) and *orco-* OSN analysis (bottom right; Fig. 3). Colored lines represent groups of nuclei, with line thickness proportional to the size of the group. Black arrowheads mark OSN clusters that were split in the subsetted analyses. (**B**) Number of *orco+*, *orco-*, *Ir76b+*, and *Ir8a+* glomeruli identified in three female brains by *Herre et al 2022* in the LVP strain of *Ae. aegypti* (grey dots)^12^ compared to the number of corresponding clusters from our analysis (red). Glomerulus counts exclude those targeted by palp neurons. snRNAseq cluster counts (red) range from the number that express a unique complement of ligand-specific receptors to the total number. Note that *Ir76b+* and *Ir8a+* cluster numbers include those identified among *orco-* OSNs (Fig. 3) plus the 2-3 identified among *orco+* OSNs (Fig. 4).

**Figure S10.**
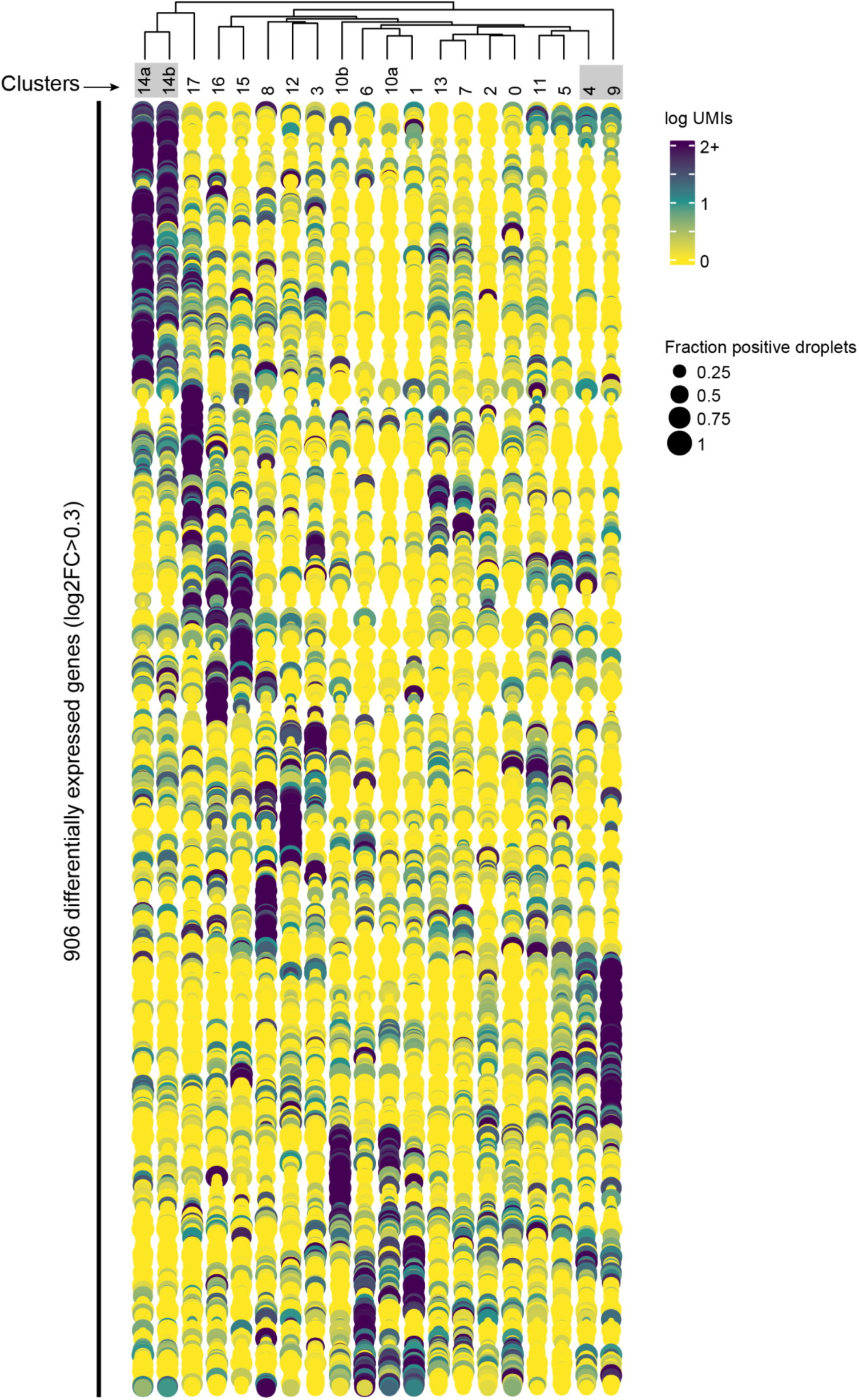
Dotplot of differential expression across 20 *orco-* OSN clusters. Grey boxes around cluster IDs highlight pairs that express the same complement of receptors and were merged in the main analysis (Fig. 3B).

**Figure S11.**
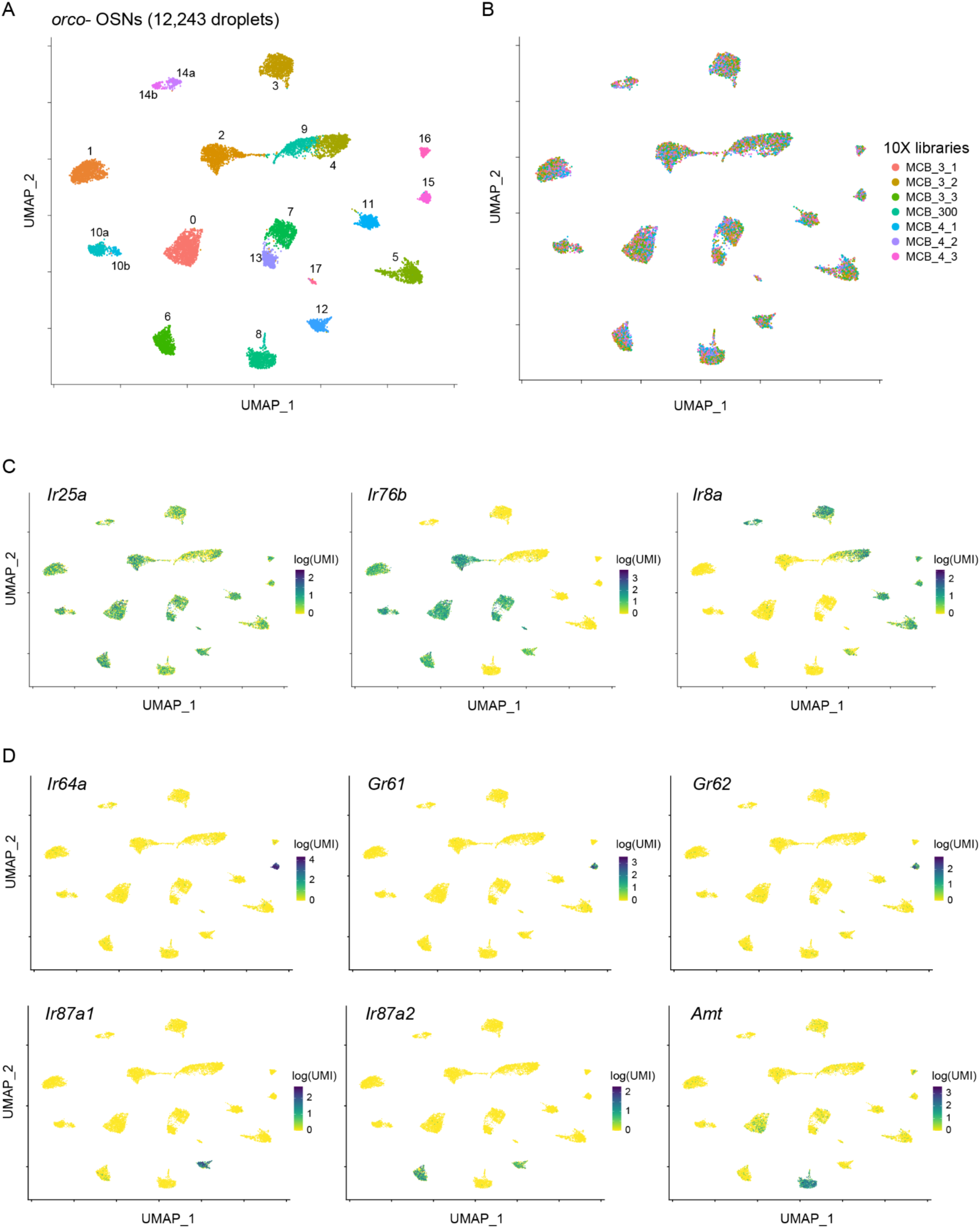
Details of UMAP clustering of *orco-* OSN subtypes. (**A**–**D**) UMAPs highlight nuclei assigned to different clusters (A), derived from different 10X libraries (B), and expressing different levels of key chemosensory co-receptors (C) or example ligand-specific receptors (D).

**Figure S12.**
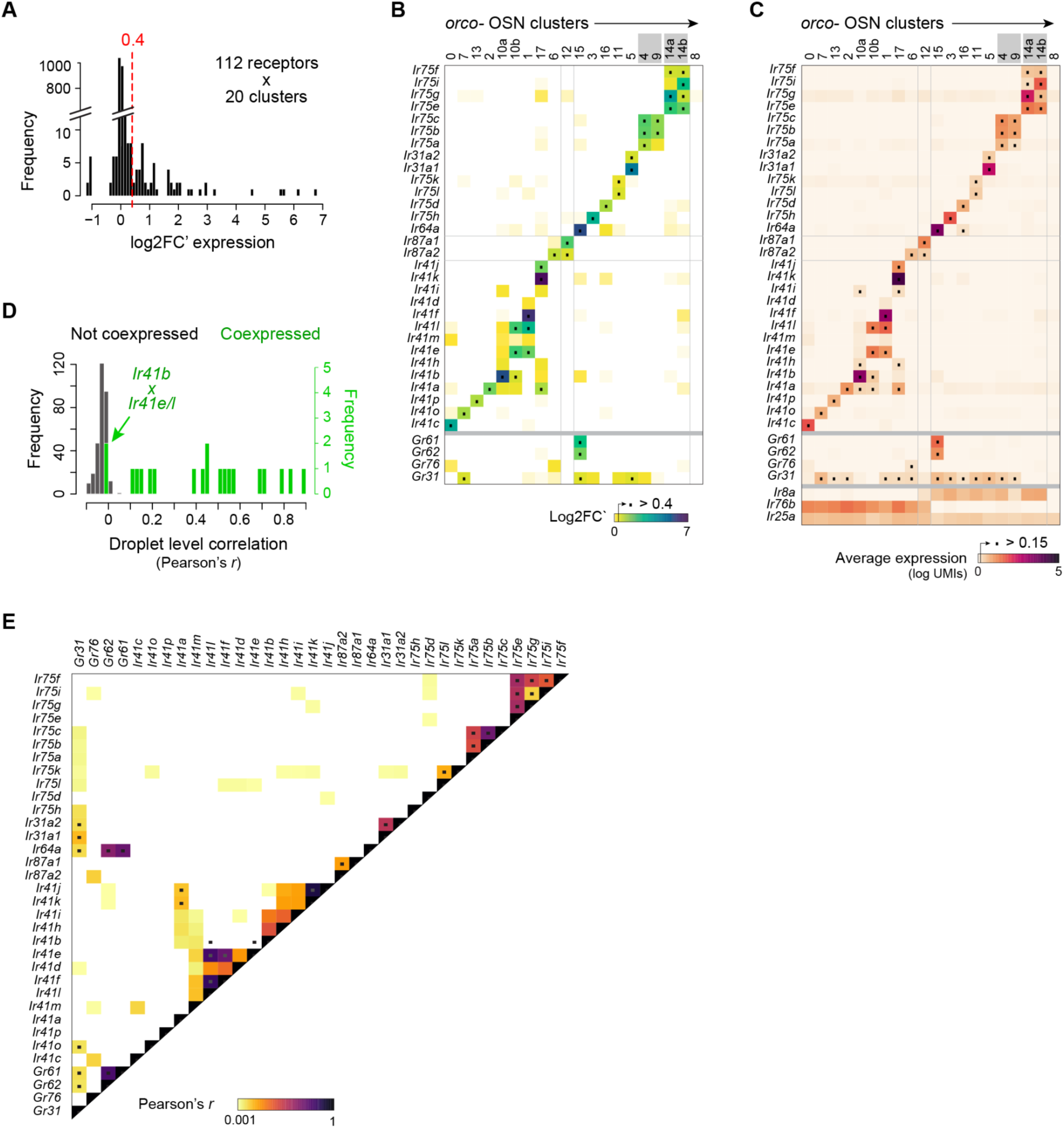
Summary of receptor expression across *orco-* OSN clusters. (**A**) Distribution of log2FC’ values used to identify a cutoff (dashed red line) for expression calls. All 112 receptors that were detected in one or more droplets were analyzed across all 20 clusters. (**B**) Heatmap showing differential expression across clusters. Plot includes all ligand-specific *IRs* (n=30), *ORs* (n=0), and *GRs* (n=4) with log2FC’ above 0.15 in any cluster, but black dots mark cases where log2FC’ exceeded the 0.4 cutoff used to call expression. Grey boxes around cluster IDs highlight pairs that express the same complement of receptors and were merged in the main analysis (Fig. 3). (**C**) Same as (B) but showing absolute expression, with black dots marking an alternative average log-scaled expression > 0.15 cutoff. (**D**) Distribution of droplet-level correlations (Pearson’s *r*) for pairs of receptors that were (red) or were not (grey) called as coexpressed based on the log2FC’ cutoff. (**E**) Pairwise droplet-level correlations for the receptors shown in (B–C). Black dots mark pairs called as coexpressed. Note that ‘coexpressed’ receptors showed elevated correlations in all but two cases. The exceptions (*Ir41b*∼*Ir41e* and *Ir41b*∼*Ir41l*) reflect a clustering artifact wherein a few *Ir41b+* droplets from cluster 10a were erroneously lumped with *Ir41e+*/*Ir41l*+ droplets in cluster 10b.

**Figure S13.**
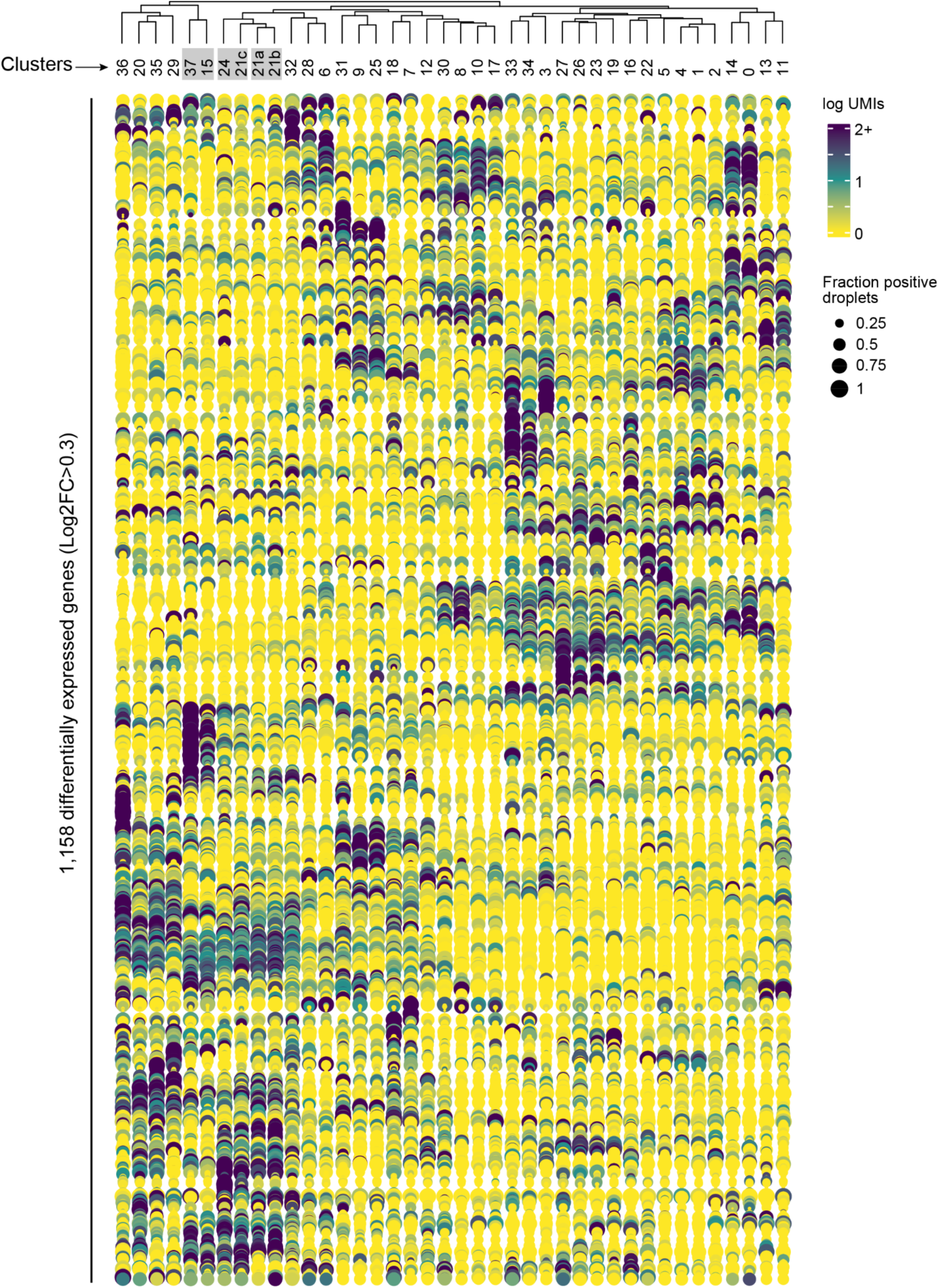
Dotplot of differential expression across 40 *orco+* OSN clusters. Grey boxes around cluster IDs highlight pairs that express the same complement of receptors and were merged in the main analysis (Fig. 4B).

**Figure S14.**
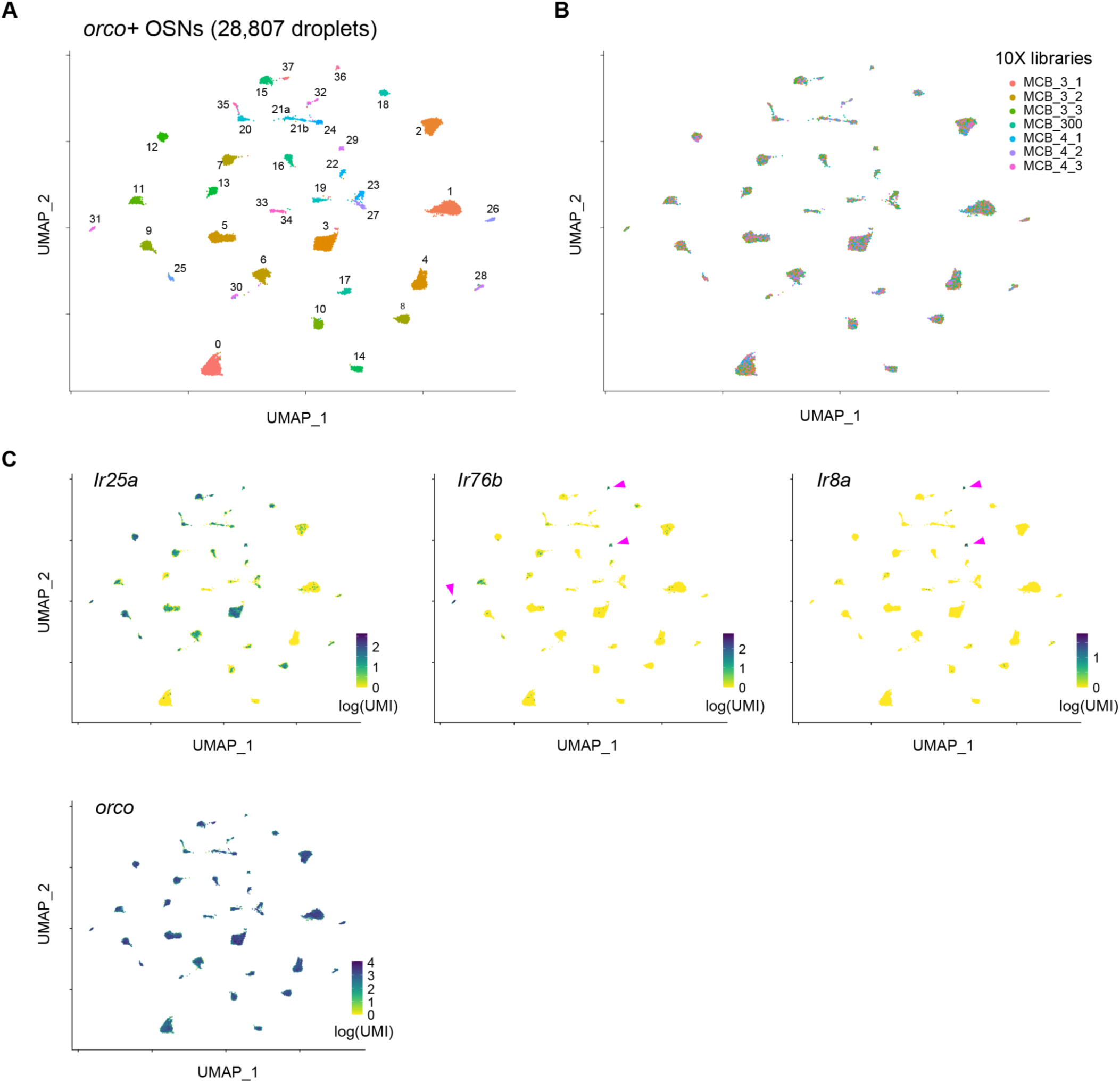
Details of UMAP clustering and co-receptor expression in *orco+* OSN subtypes. Plots highlight nuclei assigned to different clusters (**A**), derived from different 10X libraries (**B**), and expressing different levels of key chemosensory co-receptors (**C**). Pink arrow in (C) mark the few clusters showing significant *Ir76b* (n=3) and *Ir8a* (n=2) expression.

**Figure S15.**
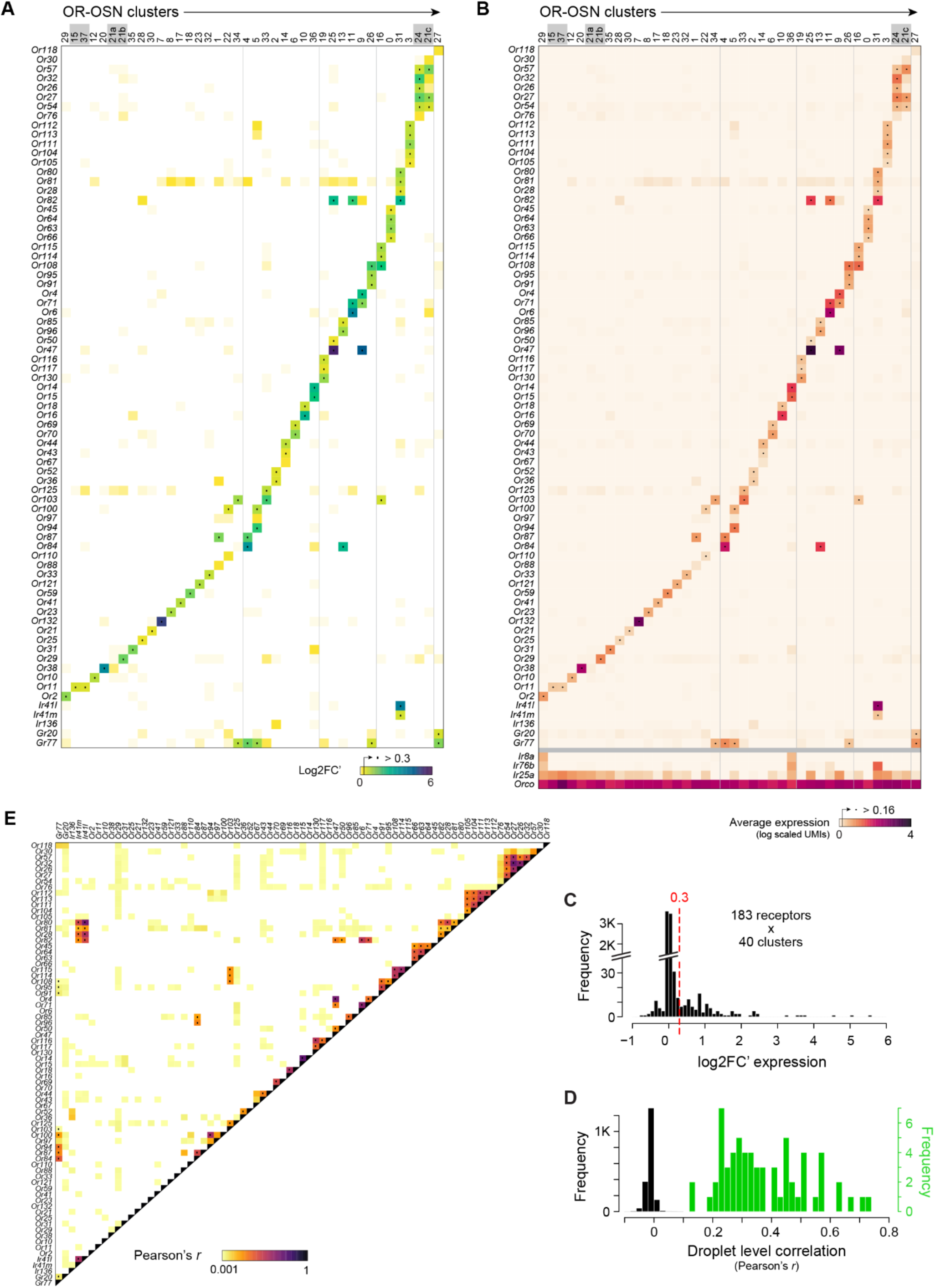
Summary of receptor expression across *orco+* OSN clusters. (**A**) Heatmap showing differential expression across clusters. Plot includes all ligand-specific receptors (*ORs*, *IRs*, *GRs*) with log2FC’ above 0.15 in any cluster, but black dots mark cases where log2FC’ exceeded the 0.3 cutoff (see C). Grey boxes around cluster IDs highlight pairs that express the same or similar complement of receptors and were merged in the main analysis (Fig. 4). (**B**) Same as (A) but showing absolute expression, with black dots marking an alternative average log-scaled expression > 0.15 cutoff.(**C**) Distribution of log2FC’ values used to identify the 0.3 cutoff (dashed red line) for expression calls. All 183 receptors that were detected in at least one or more droplets were analyzed across all 37 clusters. (**D**) Distribution of droplet-level correlations (Pearson’s *r*) for pairs of ORs that were (green) or were not (grey) coexpressed according to the log2FC’ threshold. Note the lack of overlap between the two distributions, indicating that all pairs of ORs we consider coexpressed were found not only in the same clusters, but also in the same droplets more often than expected by chance. (**E**) Pairwise droplet-level correlations for the set of receptors shown in (A–B). Black dots mark coexpressed pairs.

**Figure S16.**
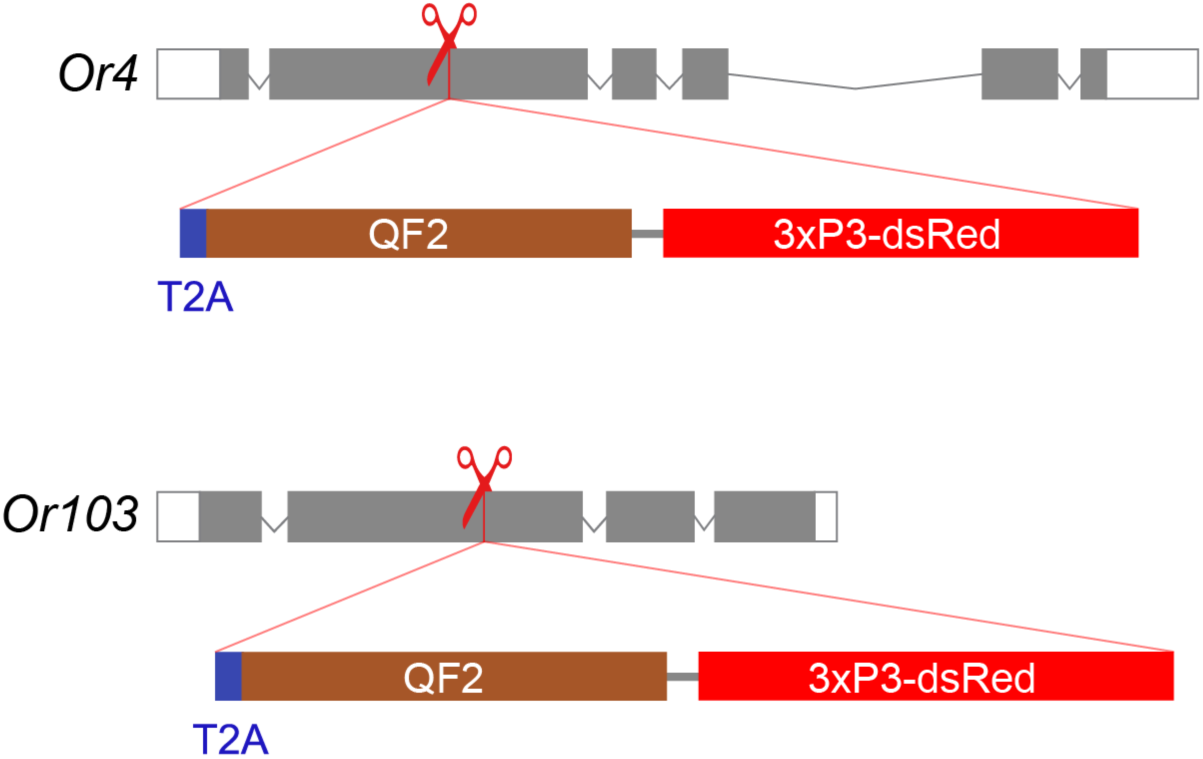
Schematic of *Or4* and *Or103* knock-in constructs. CRISPR-mediated homologous recombination was used to insert an in-frame T2A-QF2 element into the coding sequences of *Or4* and *Or103*. T2A is a ribosomal skipping sequence^66^, and QF2 is a fungal transcription factor^47^. For both loci, the insertion is expected to disrupt the coding sequence of the native receptor and result in the dual translation of the partial receptor coding sequence and QF2 from a single mRNA transcript^67^.

